# Molecular Clock Dating of Ancient Environmental DNA Reveals Damage Beyond Deamination

**DOI:** 10.64898/2026.07.03.735781

**Authors:** Maya Lemmon-Kishi, Lenore Pipes, Bianca De Sanctis, Rasmus Nielsen

## Abstract

Ancient environmental DNA (aeDNA) from permafrost, lake, cave, and marine sediments provides a rich source of genetic data that captures broad perspectives of past biodiversity. Accurate dating is crucial for discovering ecologically relevant patterns from aeDNA, and molecular clock dating would allow for sample ages to be estimated from the recovered genetic material itself instead of the geological components. However, the fragmented and damaged nature of short-read ancient DNA (aDNA) from multiple taxonomic sources poses significant challenges and has limited this dating approach for aeDNA. Here we developed ratePlacer, a phylogeny-based method for analyzing aeDNA that can combine information from many short reads in a sample while accounting for DNA damage to provide maximum likelihood estimates of sample ages. Simulations demonstrate that ratePlacer accurately dates samples even under the fragmented, damaged conditions characteristic of aeDNA and outperforms Bayesian tip-dating approaches for taxonomically mixed samples commonly found in aeDNA. Yet age estimates from re-dating Kap København varied across taxa, highlighting the difficulty of molecular clock dating in aeDNA. This dating also revealed elevated *G* → *T* and *C* → *A* mismatches consistent with oxidative damage. These patterns reveal aDNA damage beyond deamination and that remains understudied, suggesting that aeDNA should be carefully evaluated in genomic and evolutionary analyses. The new dating method, ratePlacer, extends molecular clock dating of aDNA from single-specimen to pooled environmental DNA data, where traditional methods struggle.

## 1 Introduction

Environmental DNA (eDNA) is DNA that can be extracted and sequenced from environmental sources such as soil, water, and air (Taberlet et al. 2012; Thomsen and Willerslev 2015; Clare et al. 2022; Lynggaard et al. 2022). Because all organisms shed DNA into the environment through hair, skin, feces, and blood, eDNA can reveal which organisms are present and is widely used to study ecosystems (Bohmann et al. 2014). Beyond modern sources, eDNA extracted from permafrost, caves, and lake cores, often called ancient environmental DNA (aeDNA) or sedimentary ancient DNA (sedaDNA), is used to determine the taxonomic composition of organisms that inhabited an environment in the past (Capo et al. 2021). Such analyses are possible because aeDNA is preserved for thousands to potentially millions of years by binding to minerals present in the sediment (Sand et al. 2024).

Sample age is a key input to many downstream analyses of aeDNA, such as tracking community turnover and species interactions over time (Wang et al. 2021). In aeDNA studies, sample age is estimated using non-genetic methods such as radiocarbon, optically stimulated luminescence, Uranium–Thorium, and cosmogenic nuclide burial dating (Murray and Wintle 2000; Granger and Muzikar 2001; Cheng et al. 2013; Reimer et al. 2020). Each of these approaches has specific use cases depending on the sampling location, mineral composition of the sample, and estimated age. However, these methods date the mineral or organic components of the sediment rather than the DNA, and the recovered DNA is often assumed to be contemporaneous with the dated material. This assumption is not always justified, as the processes that deposit and preserve DNA need not coincide with those dated by geological methods (Haile et al. 2007; Seeber et al. 2024). A method that estimates age from the DNA itself would not depend on this assumption.

aeDNA poses unique analytical challenges that compound those of modern eDNA and single-source ancient DNA (aDNA). Because the taxonomic origin of each read is unknown, reads must be assigned to references through competitive mapping against comprehensive databases, and ancient endogenous sequences separated from modern and microbial contamination (Pedersen et al. 2016; Wang et al. 2022). The authenticity of aDNA is established by hallmark signatures of post-mortem degradation, including short fragment length and characteristic deamination patterns (Orlando et al. 2021). In double-stranded libraries, deamination is observed as an elevated rate of *C* → *T* transitions at 5’ termini and complementary *G* → *A* transitions at 3’ termini, while deamination in single-stranded libraries is observed as elevated rates of *C* → *T* on both the 5’ and 3’ termini (Hofreiter et al. 2001; Briggs et al. 2007; Orlando et al. 2021). Furthermore, mapping accuracy relies heavily on reference database completeness and alignment parameters, while the ultra-short nature of ancient fragments inherently limits the amount of linked genetic information captured per read. There is growing interest in molecular and population genetic analyses of aeDNA beyond taxonomic identification, and damage is especially important for molecular dating. In classic molecular dating, sample ages are estimated from the substitutions that accumulate along a branch (Shapiro et al. 2011). Unaccounted damage artifacts are easily misidentified as real divergence, which artificially lengthens branches and biases age estimates (Ho et al. 2007). Most damage-aware tools are built solely around deamination, since it is the dominant and best-characterized form of post-mortem damage (Jónsson et al. 2013; De Sanctis et al. 2025b).

Deamination, however, is not the only process that degrades DNA. Oxidative damage has been measured in ancient specimens (Höss et al. 1996) and putative lesions were attributed to increases in non-deamination mismatches (Lamers et al. 2009). Early genomic-scale studies found transversions to be rare, with any slight excess of *G* → *T* attributed to low levels of oxidative damage and treated as negligible (Stiller et al. 2006; Briggs et al. 2007). Whether such damage contributes appreciably to the substitution patterns of aDNA has received little attention since, despite its relevance for analyses that depend on accurate substitution patterns, such as molecular dating.

Since aeDNA was first sequenced in 2003 (Willerslev et al. 2003), only two studies have attempted to estimate sample age from genetic information (Willerslev et al. 2007; Kjær et al. 2022), both using phylogenetic approaches to date the tips representing the aeDNA sequences. While genetic dating of individual ancient genomes is well established using the deficit of substitutions in an ancient lineage to estimate its age (Shapiro et al. 2011; Meyer et al. 2012; Prüfer et al. 2014; Valk et al. 2021), these methods remain underdeveloped for the mixed, multi-taxon assemblages typical of aeDNA. Willerslev et al. (2007) dated silty basal ice from the Dye 3 core in southern Greenland to between 450 and 800 thousand years ago, the window over which their amino acid racemization, ^10^Be/^36^Cl, luminescence, and molecular estimates were consistent. Their molecular estimate used a maximum likelihood branch-shortening analysis, constructing reference ultrametric trees for the invertebrate cytochrome oxidase subunit I (COI) amplicons and comparing the amplicon sequence branch lengths to the references. Kjær et al. (2022) later applied tip dating in BEAST (Drummond and Rambaut 2007) to ancient *Betula* and mastodon reads, supporting an age of two million years for Kap København (Kap K), following the recommendation of Shapiro et al. (2011) to estimate the age of unknown sequences in aDNA. Despite these efforts, methodological limitations continue to restrict the use of molecular dating in aeDNA studies.

The limitations differ between the previous two methods. The approach of Willerslev et al. (2007) lacks a published, reproducible pipeline and does not scale. Each query sequence was compared against a database with BLAST (Altschul et al. 1990) to gather closely related homologs, which are then aligned and placed in a phylogeny for taxonomic assignment and dating, so the analysis proceeds query by query. This was tractable for the small number of cloned amplicons in the original study but does not extend to the millions of reads produced by shotgun sequencing of aeDNA. Conversely, the consensus-based tip dating approach of Kjær et al. (2022) collapses the aeDNA reads into a single high-depth consensus sequence to represent each taxon in the phylogeny. This restricts the approach to taxa where the sample can be assumed to represent a single, genetically homogeneous population at sufficient read depth. For example, of the three plant targets identified by read depth in the Kap København samples, only *Betula* was considered to represent a single source population.

In this paper, we introduce ratePlacer, a phylogeny-based maximum likelihood molecular dating method that estimates sample age from large numbers of short reads while accounting for DNA damage, without requiring a consensus sequence or a single source population. The method is scalable to realistic sized data sets of shotgun sequencing data with millions of reads, because it stores fractional likelihoods calculated both up and down in the tree and assumes the branch lengths of the backbone tree have fixed lengths as they are estimated genome-wide. This reduces the computation to a simple one-dimensional optimization for each read within a global one-dimensional optimization for sample age. We also present a new site-specific damage and error correction method that greatly improves the molecular clock estimates.

We validate the new method on simulated data, benchmark it against tip dating in BEAST X (Baele et al. 2025), and apply it to date samples from the Kap København Formation. However, in doing so, we find mismatch patterns that deviate from canonical deamination, which both complicate molecular dating and motivate the general damage-profiling approach we incorporate into the ratePlacer pipeline. We further investigate these patterns, which are observed as elevated *G* → *T* on the 5’ termini and *C* → *A* on the 3’ termini, and putatively relate this as signatures of oxidation damage of DNA.

## 2 Results

### 2.1 ratePlacer Simulation and BEAST Comparison

To motivate ratePlacer, we first assessed whether the tip-dating approach applied by Kjær et al. (2022) recovers accurate ages under conditions typical of aeDNA. All ages are reported in substitutions per site. Using IQ-TREE Alisim (Ly-Trong et al. 2022), we simulated twenty-two modern sequences and one ancient sequence (molecular age 0.001, 125kbp), downsampled the ancient sequence to 1,000 50bp reads, and added deamination damage and sequencing error using `deamSim` from gargammel (Renaud et al. 2017) and ART (Huang et al. 2012). Reads were masked for potential deamination (*C* → *T* in the first 15 bp of the 5’ end, *G* → *A* in the last 15 bp of the 3’ end), collapsed into a consensus sequence with gaps removed from the alignment, and tip-dated with BEAST X, which dramatically underestimated the true age (mean 0.0002563; RMSE 0.0007565; Fig. 1A). Using the same simulations but taking the reads prior to damage being applied, approximating a case with sufficient read depth to correct for damage, improved the accuracy (mean 0.001082; RMSE 0.0003604; Supplementary Fig. S1B), confirming that the underestimate stemmed from damage rather than the simulation or tip-dating approach itself.

**Figure 1.**
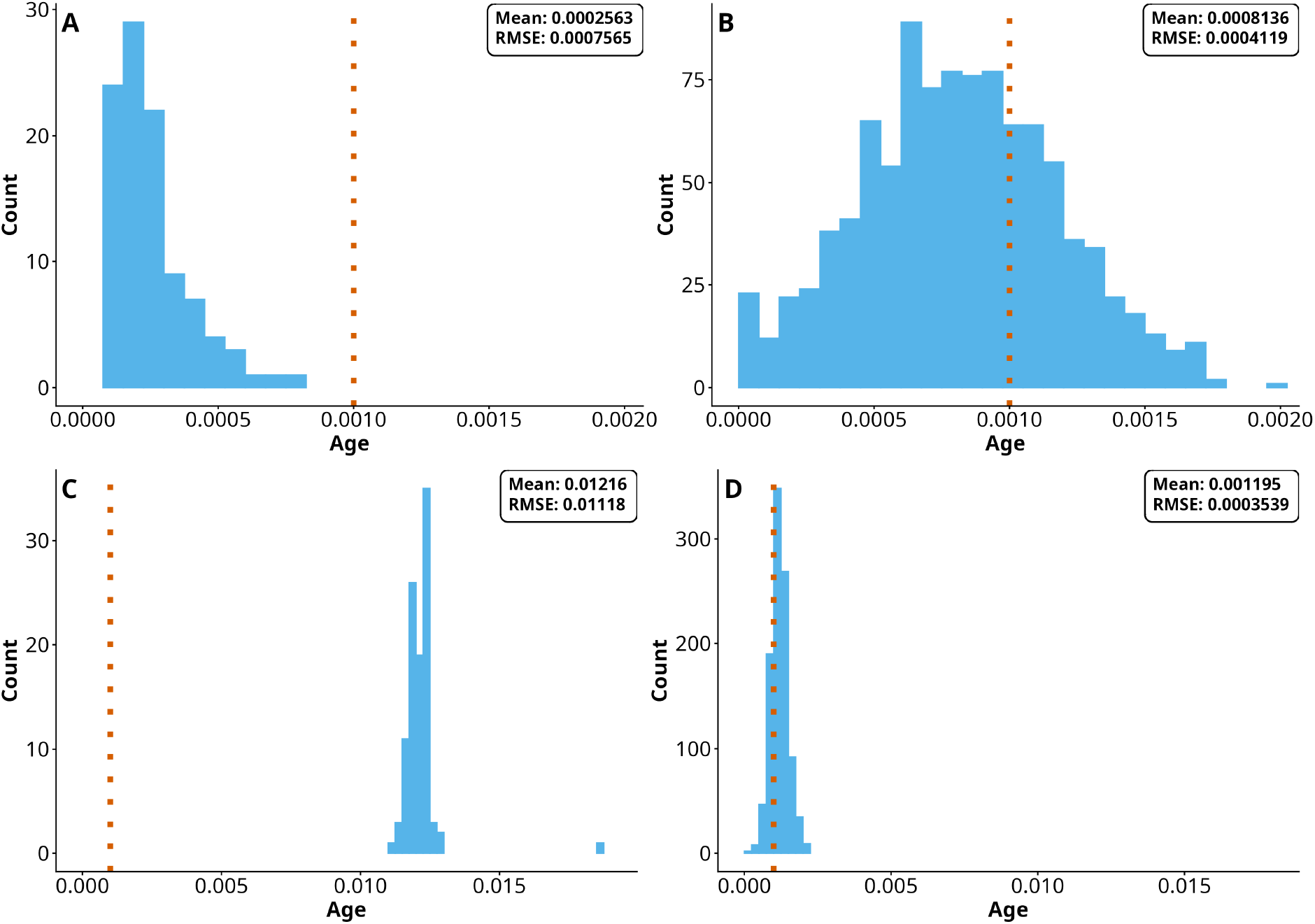
Distributions of estimated molecular ages across replicates. The dashed orange line marks the true age (0.001 substitutions/site). (A, C) BEAST X tip dating; (B, D) ratePlacer. (A, B) single-source samples; (C, D) taxonomically mixed three-taxon samples. Damage and sequencing error were added in all panels, with deamination masked at read ends. BEAST X under-estimates the age for single-source samples (A) and overestimates for mixed samples (C), whereas ratePlacer is approximately centered on the true age in both (B, D). Per-panel mean and RMSE are inset; *n* = 100 for BEAST X and *n* = 1000 for ratePlacer.

The genetic material from an environmental sample rarely derives from a single, genetically homogeneous population. To account for this, we repeated the simulation with a mixture of three ancient sequences that share a molecular age of 0.001 but represent divergent lineages within a clade. As in the previous analysis, we simulated damage and masked deamination prior to consensus generation. Despite damage biasing individual reads towards younger ages, BEAST X overestimated the age (mean 0.01216; RMSE 0.01180; Fig. 1C), likely because the consensus sequence collapses towards the ancestral sequence and loses the mutations that carry the age signal, which are unique to each species or population (Fig. 4). Entering the three ancient sequences as separate dated tips on the same simulations did not recover the true age, instead underestimating it (means 9.611 *×* 10^−5^, 3.762 *×* 10^−5^, and 6.240 *×* 10^−5^; Supplementary Fig. S1C). Two effects combine here: damage biases each ancient sequence toward younger ages (seen above in the single ancient sequence case), and because coverage is low across three taxa, aligning the reads to the reference leaves each ancient sequence with extensive missing data, which further distorts the branch estimates. Splitting the taxa therefore avoids the consensus collapse seen in Fig. 1C but introduces its own downward bias. Thus, under realistic aeDNA conditions, BEAST X tip dating performs poorly when damage and sequencing error cannot be fully corrected (e.g., low coverage) or when samples are taxonomically mixed (e.g., genus-level or higher classification).

On the same simulations, ratePlacer recovered ages with less bias. To show this, we assigned each read to a node with tronko (Pipes and Nielsen 2024), then determined its final placement by maximum likelihood (ML) across the candidate branches surrounding that node. In addition to masking potential deamination in the first and last 15 bases, we incorporated base quality scores into the per-base nucleotide likelihoods. With this correction, ratePlacer’s mean estimated age was 0.0008136 (RMSE 0.0004119; Fig. 1B). Without damage correction, the mean estimate collapsed to approximately 0 (3.247 *×* 10^−8^; RMSE 0.001), which is the minimum value ratePlacer can return, again reflecting branch lengthening from uncorrected damage alone rather than a meaningful age.

The source of this residual bias after damage correction was localized through a series of simulations (Supplementary Fig. S2G-J): with true placement, the downsampled estimate was unbiased (mean 0.0009833; G), introducing tronko placement alone shifted the mean modestly (mean 0.0008042; H), uncorrected damage drove the mean to 0 as described above (I), and the deamination damage correction restored most of the signal back to that of the tronko placement (mean 0.0008136; J). The direction of this bias is tree-dependent and not always younger (Supplementary Fig. S2). For taxonomically mixed samples, ratePlacer again recovered the age more accurately (mean 0.001195; RMSE 0.0003539; Fig. 1D; Supplementary Fig. S2L-O).

ratePlacer remained robust across molecular ages. Repeating the damage corrected, tronko-assigned, ML-placed simulations for ancient sequences at ages 0.01 and 0.0001 produced mean estimates of 0.009819 and 9.802*×*10^−5^ (RMSE 0.0004917 and 5.138*×*10^−5^, respectively; Supplementary Fig. S2E and T).

### 2.2. DNA Damage Patterns

aDNA studies typically only consider deamination damage, which presents as single base pair mismatches (*C* → *T* and *G* → *A*) at fragment ends. This damage is routinely addressed in downstream analyses by trimming or masking the terminal bases of each read. While applying ratePlacer to the double-stranded Kap København samples (Kjær et al. 2022), we observed mismatch patterns inconsistent with deamination-only damage even after end masking (Section 2.3). To characterize these patterns and assess whether they represent a broader feature of aeDNA, we performed a systematic mismatch analysis on the four most abundant genera by total nuclear read counts from Kap København, and on two additional samples differing in library preparation and preservation context. The decomposition framework introduced in Section 4.3 is motivated by the patterns described here. Following the competitive mapping procedure in Section 4.5.1, 10,301,910 *Betula*, 13,864,022 *Populus*, 23,894,466 *Salix*, and 14,495,097 *Dryas* reads were assigned to their respective nuclear genomes, covering 456,976,328, 647,211,983, 1,115,195,471, and 630,574,737 bases respectively. Observed mismatch counts and observed-to-expected ratios are shown in Table 1. Expected substitution counts are computed per taxon under a GTR+Γ model fit to the reference phylogeny (Section 4.2.2), such that observed-to-expected ratios should be equal among substitution types in the absence of damage. To evaluate whether the reversibility assumption of GTR affected these ratios, we recalculated expected counts under a non-reversible (UNREST) model; this did not change the interpretation of the results (Supplementary Section S2.1).

**Table 1.** Observed mismatches and observed-to-expected ratios of most abundant genera in Kap København. Observed mismatches and observed-to-expected ratios of *Betula, Salix, Populus*, and *Dryas* nuclear assigned reads from the Kap København samples. Observed-to-expected ratios are scaled such that the smallest ratio is 1.00.

| Substitution | *Betula* |  | *Salix* |  | *Populus* |  | *Dryas* |  |
| --- | --- | --- | --- | --- | --- | --- | --- | --- |
|  | Obs. | Ratio | Obs. | Ratio | Obs. | Ratio | Obs. | Ratio |
| $A \rightarrow C$ | 273,469 | 1.000 | 870,289 | 1.060 | 419,970 | 1.022 | 229,779 | 1.000 |
| $C \rightarrow A$ | 411,717 | 1.506 | 1,124,076 | 1.382 | 540,696 | 1.358 | 457,288 | 1.990 |
| $A \rightarrow G$ | 918,710 | 1.225 | 2,347,649 | 1.010 | 1,325,018 | 1.098 | 678,472 | 1.073 |
| $G \rightarrow A$ | 2,824,100 | 3.767 | 5,830,100 | 2.574 | 3,289,481 | 2.797 | 4,213,373 | 6.664 |
| $A \rightarrow T$ | 455,097 | 1.011 | 1,688,303 | 1.037 | 779,240 | 1.005 | 385,589 | 1.063 |
| $T \rightarrow A$ | 454,674 | 1.010 | 1,667,899 | 1.026 | 771,639 | 1.000 | 387,036 | 1.067 |
| $C \rightarrow G$ | 175,113 | 1.109 | 446,074 | 1.244 | 209,741 | 1.174 | 145,295 | 1.055 |
| $G \rightarrow C$ | 173,929 | 1.101 | 444,008 | 1.242 | 209,225 | 1.170 | 144,584 | 1.050 |
| $C \rightarrow T$ | 2,998,741 | 3.977 | 6,087,380 | 2.660 | 3,465,224 | 2.924 | 4,572,011 | 7.172 |
| $T \rightarrow C$ | 914,849 | 1.213 | 2,338,905 | 1.000 | 1,324,176 | 1.092 | 670,941 | 1.053 |
| $G \rightarrow T$ | 414,848 | 1.500 | 1,134,071 | 1.380 | 543,193 | 1.351 | 467,420 | 2.015 |
| $T \rightarrow G$ | 279,091 | 1.009 | 879,512 | 1.064 | 426,192 | 1.034 | 240,803 | 1.038 |

As expected, deamination-associated substitutions (*C* → *T* and *G* → *A*) violated symmetry across all four taxa (3.278, 2.603, 2.617, and 6.814 fold for *C* ↔ *T*; 3.074, 2.483, 2.483, and 6.210 fold for *G* ↔ *A*) and showed the highest observed-to-expected ratios within each taxon. Symmetry was also violated for *G* → *T* and *C* → *A* (1.486, 1.289, 1.275, and 1.941 fold for *G* ↔ *T*; 1.506, 1.292, 1.288, and 1.990 fold for *C* ↔ *A*), while *A* ↔ *T* and *C* ↔ *G* remained symmetric (0.996–1.012 and 1.003–1.007 fold respectively). *G* → *T* and *C* → *A* were consistently the third and fourth highest observed-to-expected ratios across all four taxa; the ranks of the remaining substitution types showed no discernible pattern. *G* → *T* and *C* → *A* are canonical signatures of oxidative damage from 8-oxoguanine misincorporation (Shibutani et al. 1991; Costello et al. 2013).

Positional substitution patterns along the read were examined for *Betula* (Fig. 2; remaining taxa in Supplementary Fig. S3-S5). Deamination damage and non-damage-associated substitutions are plotted separately (panels A,C vs. B,D) to visualize them on comparable scales. As expected, *C* → *T* rates were elevated at the 5’ end and *G* → *A* at the 3’ end, decaying to an elevated background rate within roughly eight bases of the termini, with observed-to-expected following a similar pattern and the ratios reaching 32.230 and 30.574 at the terminal positions. *G* → *T* and *C* → *A* were also elevated at the read ends, with *G* → *T* reaching 4.098 at the 5’ end and *C* → *A* reaching 3.798 at the 3’ end. The asymmetry between these substitutions across the two ends is expected for double-stranded libraries as *G* → *T* and *C* → *A* are complementary readouts of the same underlying damage from opposite strands. Like deamination, both substitutions remained elevated above their complements (*T* → *G* and *A* → *C*) across the read interior in both observed rates and observed-to-expected ratios. *C* ↔ *G* was uniformly low and symmetric at every position. *A* → *T* and *T* → *A* were symmetric across the read interior but violated symmetry at the terminal ends; *A* → *T* reached an observed-to-expected ratio of 4.773 at position 1 of the 5’ end and *T* → *A* reached 4.738 at position −1 of the 3’ end. Unlike the damage substitutions, this elevation was confined to a single terminal position with opposing directionality at each end, which is inconsistent with damage chemistry and more consistent with a terminal-base artifact from library preparation or mapping. Both deamination and oxidation damage signatures persist across the read interior beyond the terminal regions where they are most pronounced, indicating that neither is restricted to fragment ends. This persistence extended well beyond position 10: at position 15 of the 5’ end *C* → *T* and *G* → *T* retained observed-to-expected ratios of 2.189 and 1.319, respectively, and at position −15 of the 3’ end *G* → *A* and *C* → *A* retained ratios of 2.186 and 1.311, respectively. The same qualitative patterns held in the other three taxa (Supplementary Fig. S3-S5).

**Figure 2.**
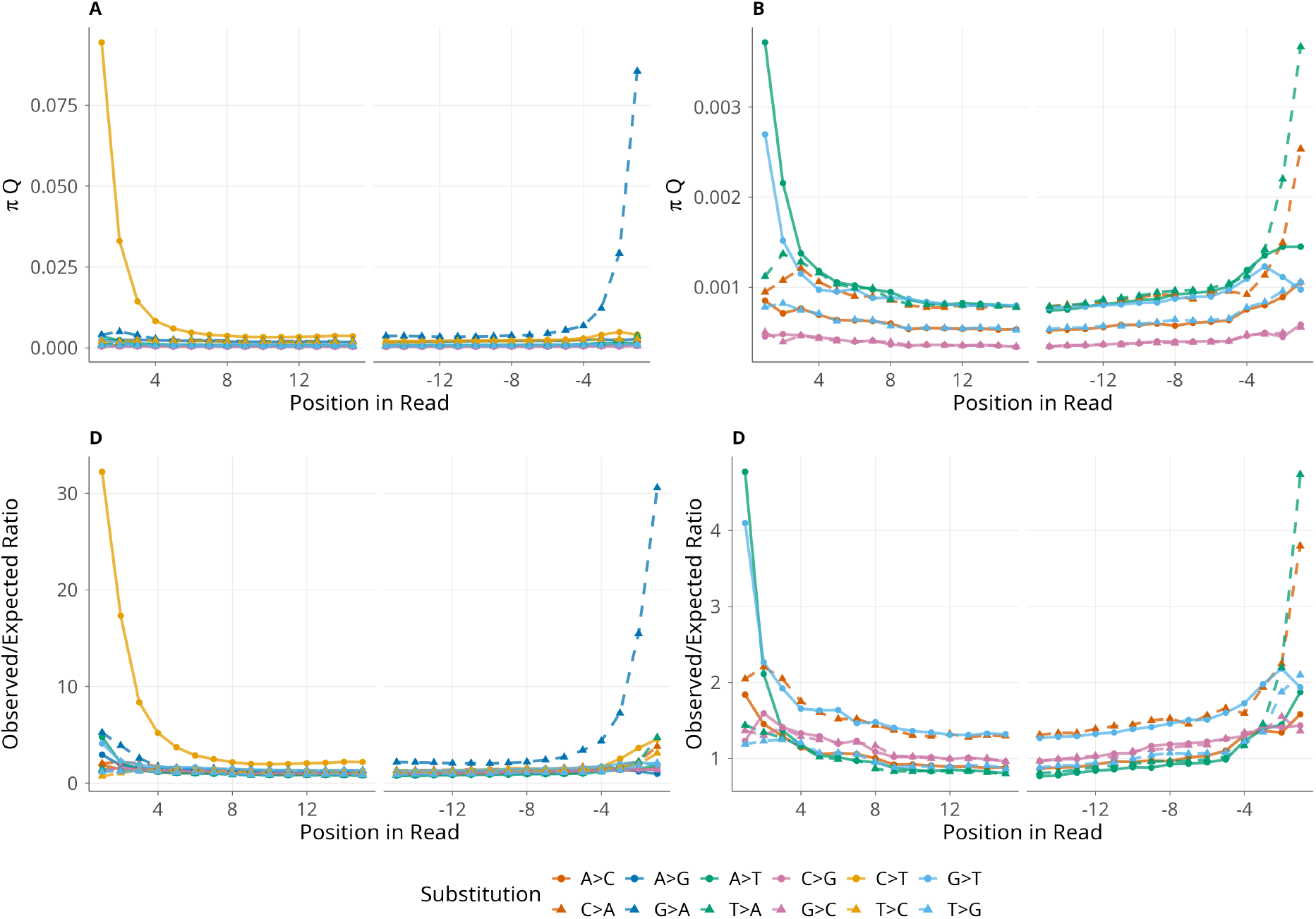
Positional substitution patterns for *Betula* reads at the first and last fifteen positions. Panels (A) and (C) show all substitutions; panels (B) and (D) show non-deamination damage substitutions (*C* → *T, G* → *A*, and symmetric pairs are removed). Deamination damage substitutions are removed from panels (B) and (D) due to differences in scale. Panels (A,B) show observed rates (*πQ*); panels (C,D) show observed-to-expected ratios, with the same scaling used for *Betula* in Table 1. Solid lines denote one direction of each substitution pair and dashed lines denote the reverse (e.g., solid *C* → *T* vs. dashed *T* → *C*). Under the symmetry assumption, matched dashed and solid lines of the same color should overlap in (A,B), and all lines should overlap in (C,D); gaps between matched pairs indicate violations of symmetry. Elevation above the background for both deamination (*C* → *T, G* → *A*) and oxidative damage (*G* → *T, C* → *A*) substitutions is visible across the interior of the read in addition to the terminal positions.

To determine whether *G* → *T* and *C* → *A* are elevated specifically in ancient reads or all reads, we calculated the symmetry ratio for each substitution type on 192,258,396 ancient and 5,404 modern reads (Fig. 3A). Consistent with the per-taxon results, ancient reads showed elevated *C* → *T* and *G* → *A* (5.7402 and 5.4297), as well as *G* → *T* and *C* → *A* (1.9239 and 1.9785), while *A* → *T* and *C* → *G* remained symmetric (0.9963 and 1.0021). Modern reads showed no comparable excess (1.1213, 1.2958, 1.0732, 0.8788, 1.2510, and 1.1008 for *C* → *A, G* → *A, A* → *T, C* → *G, C* → *T*, and *G* → *T* respectively). *χ*^2^ tests with Cramér’s V effect sizes (Table 2) found all symmetric pairs significant except for *A* ↔ *T*. The *C* ↔ *G* pair was significant at *p* = 0.0070, while the remaining four pairs yielded *p <* 1.0 *×* 10^−10^. All Cramér’s V values were small (*<* 0.01), indicating that the proportional deviations from symmetry are modest in magnitude. The strong *χ*^2^ significance is driven by the very large sample sizes (*>* 192M ancient reads), which detect even small deviations as significant. Despite the small Cramér’s V values, the per-substitution counts (Table 1, all *>* 10^5^) are sufficient to resolve relative ordering of effect sizes with high confidence; the ranking of damage-associated pairs above non-damage pairs is therefore unlikely to reflect sampling variation. The relative ranking of Cramér’s V across substitution pairs places the damage-associated pairs (*C* ↔ *T* = 0.0060, *G* ↔ *A* = 0.0057, *C* ↔ *A* = 0.0038, *G* ↔ *T* = 0.0036), well above *C* ↔ *G* (0.0013) and *A* ↔ *T* (0.0003).

**Table 2.** *χ*^2^ tests and Cramér’s V effect sizes comparing ancient and modern substitution ratios at Kap København. Symmetric pairs test whether the rate of each substitution equals its reverse (e.g., *C* → *T* vs. *T* → *C*); complement pairs test whether substitutions on opposite strands occur at equal rates under Watson-Crick base pairing (e.g., *A* → *C* vs. *T* → *G*).

| Substitution pair | Ancient ratio | Modern ratio | P-value | Cramér’s V |
| --- | --- | --- | --- | --- |
| *Symmetric pairs* (substitution and its reverse) |  |  |  |  |
| $C \rightarrow T / T \rightarrow C$ | 5.7402 | 1.2510 | $<1 \times 10^{-10}$ | 0.0060 |
| $G \rightarrow A / A \rightarrow G$ | 5.4297 | 1.2958 | $<1 \times 10^{-10}$ | 0.0057 |
| $C \rightarrow A / A \rightarrow C$ | 1.9785 | 1.1213 | $<1 \times 10^{-10}$ | 0.0038 |
| $G \rightarrow T / T \rightarrow G$ | 1.9239 | 1.1009 | $<1 \times 10^{-10}$ | 0.0036 |
| $C \rightarrow G / G \rightarrow C$ | 1.0021 | 0.8788 | $7.0 \times 10^{-3}$ | 0.0013 |
| $A \rightarrow T / T \rightarrow A$ | 0.9963 | 1.0732 | $2.2 \times 10^{-1}$ | 0.0003 |
| *Complement pairs* (Watson-Crick complements) |  |  |  |  |
| $C \rightarrow G / G \rightarrow C$ | 1.0021 | 0.8788 | $7.0 \times 10^{-3}$ | 0.0013 |
| $A \rightarrow C / T \rightarrow G$ | 0.9643 | 1.0421 | $1.3 \times 10^{-1}$ | 0.0006 |
| $G \rightarrow T / C \rightarrow A$ | 1.0084 | 0.9421 | $1.6 \times 10^{-1}$ | 0.0004 |
| $A \rightarrow T / T \rightarrow A$ | 0.9963 | 1.0732 | $2.2 \times 10^{-1}$ | 0.0003 |
| $A \rightarrow G / T \rightarrow C$ | 1.0037 | 0.9747 | $4.5 \times 10^{-1}$ | 0.0002 |
| $C \rightarrow T / G \rightarrow A$ | 1.0533 | 0.9905 | $6.1 \times 10^{-2}$ | 0.0002 |

**Figure 3.**
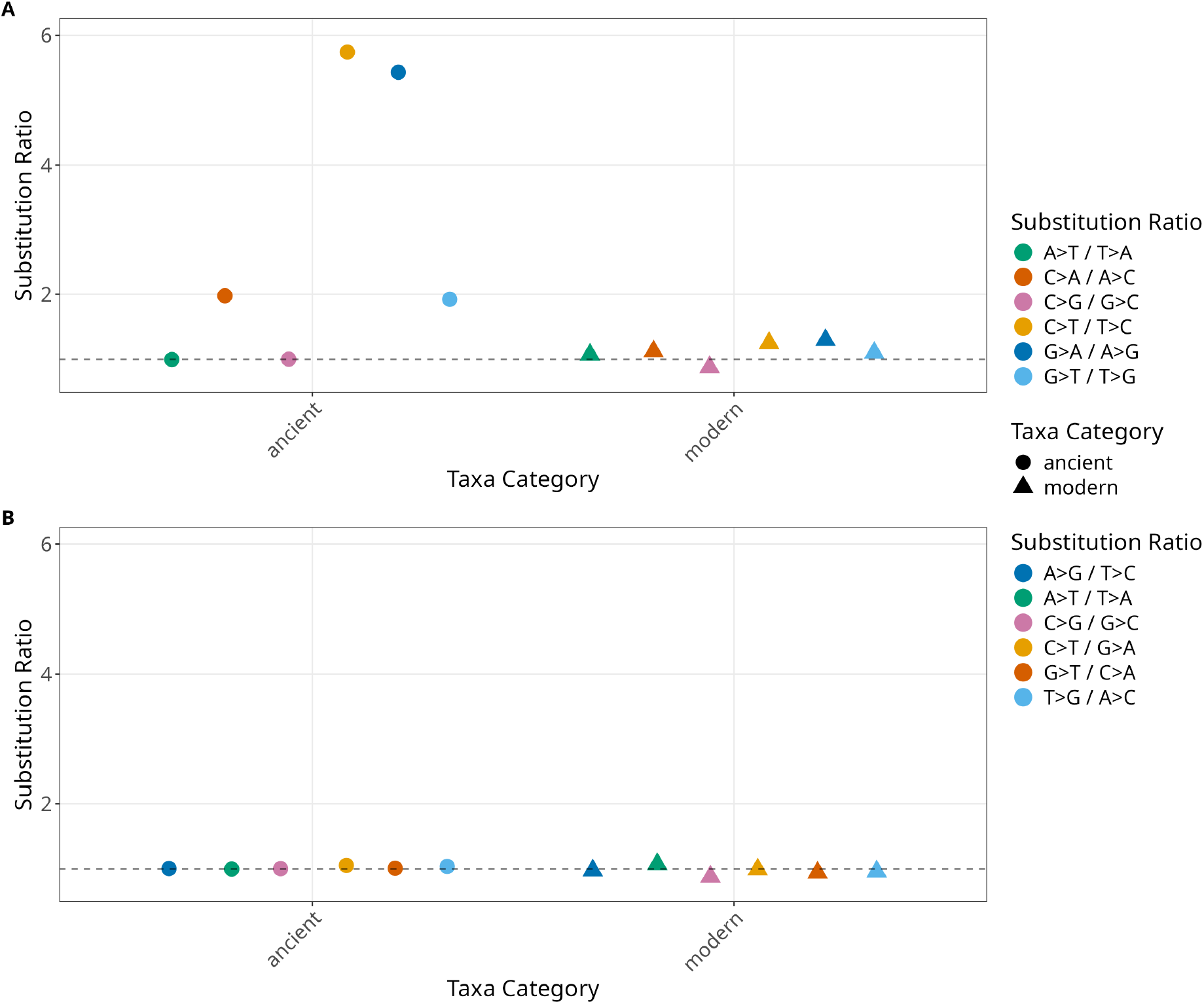
Substitution symmetry ratios for ancient versus modern read assignments at Kap København. Each point is one substitution ratio; box plots summarize the distribution across all six ratios within a category. The dashed line marks the expected ratio of 1.0 under the relevant null. (A) Symmetric substitution ratios (*i* → *j* / *j* → *i*), which should equal 1.0 under time-reversible substitution. Ancient reads show elevated ratios for the four damage-associated pairs (*C* → *T* /*T* → *C, G* → *A*/*A* → *G, C* → *A*/*A* → *C, G* → *T* /*T* → *G*). (B) Watson-Crick complement ratios, which should equal 1.0 in double-stranded libraries where strand of origin is unknown. Both ancient and modern reads remain near 1.0, ruling out strand-specific sequencing bias as the source of the symmetry violations in (A).

Sequencing error and mapping misalignment were ruled out by examining Watson-Crick complement pairs (substitution complement pairs such as *C* → *T/G* → *A* and *G* → *T/C* → *A*), which ranged from 0.8788 to 1.0732 in both ancient and modern reads (Table 2, complement pairs; Fig. 3B). All the Watson-Crick pairs were insignificant in the *χ*^2^ test with the exception of *C* → *G/G* → *C*, which was significant at *p* = 0.0070. The effect of taxonomic assignment was tested by permuting ancient/modern labels 10,000 times (Section 4.2.3). The observed ancient-modern difference was significant for *C* → *T, G* → *A, G* → *T*, and *C* → *A* (*p <* 1.0 *×* 10^−10^, *p <* 1.0 *×* 10^−10^, 0.0040, and 0.0042 respectively) and non-significant for *A* → *T* and *C* → *G* (Supplementary Fig. S6-S7). Combined with the symmetric Watson-Crick ratios, this indicates that the observed excess of *G* → *T* and *C* → *A* likely reflects DNA misincorporation from environmental oxidative damage rather than laboratory or pipeline bias.

To test whether oxidative damage signatures are unique to Kap København, we applied the symmetry analysis to two additional libraries from other sources: Krestovka mammoth molar (Valk et al. 2021) and subglacial Antarctic precipitates (De Sanctis et al. 2025a). The Krestovka mammoth molar is a Uracil-DNA Glycosylase (UDG) treated double-stranded library from ancient bone DNA, dated to the Early Pleistocene and found in the Lower Olyorian deposits. In the 1,123,009 nuclear reads that mapped to *Elephas maximus*, only *G* → *T* and *C* → *A* violated symmetry at 2.7299 and 3.2118 compared to other substitution types ranging from 0.8661 to 1.1502 (Table 3). In contrast, the subglacial Antarctic precipitate samples are single-stranded libraries from Antarctica and date between 16,000 and 570,000 years ago. This sample showed violations only for deamination-associated *C* → *T* with a ratio of 2.7875 compared to the rest that ranged from 1.0339 to 1.1133.

**Table 3.** Symmetry ratios across samples differing in library preparation and preservation context. Ratios are computed as max(*i* → *j, j* → *i*)*/* min(*i* → *j, j* → *i*), so all values are ≥ 1.0; values near 1.0 indicate symmetry. Kap København modern reads were filtered against control-sample contamination (Supplementary Section S2.2). Only reads from subglacial precipitates were used (Supplementary Section S2.5).

| Sample | Library | Treatment | $C \leftrightarrow T$ | $G \leftrightarrow A$ | $C \leftrightarrow A$ | $G \leftrightarrow T$ | $A \leftrightarrow T$ | $C \leftrightarrow G$ |
| --- | --- | --- | --- | --- | --- | --- | --- | --- |
| Kap K (ancient) | ds | — | 5.7402 | 5.4297 | 1.9785 | 1.9239 | 1.0037 | 1.0021 |
| Kap K (modern) | ds | — | 1.2510 | 1.2958 | 1.1213 | 1.1009 | 1.0732 | 1.1379 |
| Krestovka mammoth | ds | UDG | 1.1405 | 1.1502 | 3.2118 | 2.7299 | 0.8661 | 1.0210 |
| Antarctic (subglacial) | ss | — | 2.7875 | 1.1133 | 1.0781 | 1.0939 | 1.0440 | 1.0339 |

The oxidative damage signature observed at Kap København can therefore also be found in aDNA from bones. This is consistent with an in situ origin for the oxidative damage, though additional samples spanning a wider range of preservation environments would be needed to characterize its environmental dependence.

### 2.3. Re-estimation of the Molecular Age of Kap København

To validate ratePlacer, we first estimated the age of Kap København using the same Betula chloroplast consensus sequence generated by Kjær et al. (2022). From this consensus *Betula* ancient sequence and the aligned chloroplast references from the BEAST dating, the estimated molecular age was 0.0004049 expected substitutions per site. Using the *Betula-Alnus* divergence of 61.1 Ma from the Kap K BEAST dating (Yang et al. 2019), this molecular age was scaled against the ultrametric reference tree (depth 0.01516 substitutions per site), yielding an age of ~1.631 Ma. This is older than the median age of 1.323 Ma reported by Kjær et al. (2022), but falls within their 95% highest posterior density (HPD) of 0.6786 to 2.0172 Ma. The 95% confidence interval of the ratePlacer estimate ranges from 1.631 to 1.632 Ma (molecular ages of 0.0004047-0.0004050). The two intervals capture different sources of uncertainty and are not directly comparable, as the ratePlacer interval conditions on a fixed reference tree, divergence time, and substitution model, whereas the Kjær et al. (2022) highest posterior density interval propagates uncertainty in these jointly with the clock rate and topology.

We next attempted to estimate the age of *Betula* directly using the mapped nuclear reads (*N* = 10, 301, 840 reads), masking deamination at the first and last 7 bases (as the corresponding deamination plot indicates damage plateaus by this point). Using ML-based read placement, the age estimate was 0.000000032986, very near the smallest age ratePlacer can return. We also estimated the age by masking all possible transitions and trimming the ends by 7 bp, but neither treatment significantly changed the age estimate. The estimates returned ages near the smallest value ratePlacer can produce suggested unaccounted damage as in simulations, this floor was reached only when no damage correction was applied. This also explains the contrast with the chloroplast consensus above, which was built from sites of at least 20x depth and effectively masked both deamination and oxidative damage. The nuclear age estimate has no such protection. Placing reads individually exposes the interior deamination and oxidative damage that the consensus had removed.

Damage asymmetry analysis subsequently showed that damage was more complex than terminal deamination: potential oxidative damage was also present and both were elevated across the full read length. We therefore applied the damage decomposition method to estimate damage and recalculate nucleotide likelihoods (4.3). Throughout, we report two estimates per taxon: *α*_*hi*_ and *α*_*lo*_, corresponding to high and low values of the evolutionary-time parameter *α*, where *α*_*lo*_ attributes a greater share of observed mismatch to damage. Using this damage correction, we obtained molecular ages of 0.001504 (0.001498-0.001510, *α*_*hi*_) and 0.001451 (0.001444-0.001458, *α*_*lo*_). The molecular clock used a *Betula-Alnus* divergence ranging from ~58 Ma (Yang et al. 2025) to ~61 Ma (Yang et al. 2019), with an ultrametric reference tree depth of 0.09584. This gives a possible time range for Kap København of 880 ka to 5.33 Ma from *Betula* dating (point estimates ranging ~884 to 963 ka).

We applied the same approach to *Populus, Salix*, and *Dryas*, the other most abundant ancient genera. For *Populus* (*N* = 13, 826, 034 reads), ratePlacer estimated molecular ages of 0.0097021 (0.009686-0.0097183, *α*_*hi*_) and 0.01136 (0.01134-0.01137, *α*_*lo*_). Divergence estimates for *Populus-Salix* vary from ~35 Ma (Sanderson et al. 2023) to ~52 Ma (Dai et al. 2014); with a tree depth of 0.1394, this converts to ~2.44 to ~4.24 Ma, the maximum likelihood estimates falling in the same range. For *Salix* (N = 23,747,639 reads), ratePlacer returned molecular ages of 0.01052 (0.01051-0.01054, *α*_*hi*_) and 0.01207 (0.01205-0.01208, *α*_*lo*_). Using the same divergence times and a tree depth of 0.1247, this converts to ~2.95-5.04 Ma, the point estimates spanning the same range. Finally, for *Dryas* (N = 14,495,016 reads) we obtained molecular ages of 0.001977 (0.001968-0.001985, *α*_*hi*_) and 0.002027 (0.002019-0.002035, *α*_*lo*_). The *Dryas-Purshia* divergence is estimated at ~38 Ma (Xiang et al. 2017) to 40.67 Ma (Zhang et al. 2017); with a tree depth of 0.06427, this converts to ~1.16-1.29 Ma. Despite the inherent uncertainty in genus-level divergence times, the molecular ages for all three taxa converge on a Plio-Pleistocene origin. These independent estimates are broadly consistent with each other and align with the accepted two-million-year age of the site.

## 3. Discussion

In this paper we introduced ratePlacer, a molecular dating method developed for aeDNA. Across simulations, ratePlacer estimates age more accurately than tip dating with BEAST X, as it requires neither a consensus sequence nor a single source population. Applied to Kap København, ratePlacer revealed evidence of oxidative damage of guanine and motivated a new approach to profiling aDNA damage. Even with this correction, age estimates across taxa did not fully agree, underscoring the difficulty of molecular dating from aeDNA. Nonetheless, the estimates remain compatible with the original claims that the DNA from Kap København is two million years old, and possibly older.

Our simulations show that ratePlacer recovers accurate molecular ages across a wide temporal range and under the deamination damage and mixed taxa assemblages typical of aeDNA, and does so more reliably than consensus based tip dating in BEAST X. These results held under a more stringent test than the empirical Kap K analysis, with far lower coverage and so less signal for age estimation. At low read depth, even when read ends are masked for deamination, the consensus approach underestimates age, likely because of deamination at internal positions of the read, and we expect the oxidative damage uncovered in the Kap K reanalysis to exacerbate this. It fails further when multiple species are present, because collapsing reads into a single sequence masks evolutionary substitutions and draws the sequence toward the ancestral state. Even in idealized scenarios where reads are perfectly separated by species, jointly estimating their ages biases the estimates younger. Uneven read coverage introduces gaps into the multiple sequence alignment that inflate the branch lengths of the ancient sequences, and restricting the estimate to conserved regions is not possible because competitive mapping is biased toward assigning reads to less conserved regions. Although we did not test this directly, we caution against the consensus approach even when a single species is confidently identified, because collapsing a population discards genetic diversity in the same way (Fig. 4). This effect matters far more for young samples, where the lost diversity is a larger fraction of the branch, than for old ones such as Kap K.

Initial application of ratePlacer to the Kap København dataset failed to yield a meaningful age, as the program returned its computational lower bound. In simulation this behavior is diagnostic of uncorrected damage, and recognizing it led us to identify putative oxidative damage of guanine in these samples. From sequencing data alone we cannot assign a mechanism to the oxidative modification of guanine, though Höss et al. (1996) previously found 8-hydroxyguanine in all of their samples. Oxidative damage of guanine is well characterized in other fields such as cancer genomics, where it produces *G* → *T* substitutions and complementary *C* → *A* (Costello et al. 2013). As with deamination, the mismatch arises from misincorporation opposite the modified guanine. The observed *G* → *T* rate is a lower bound on the true extent of oxidative damage in our samples, because this misincorporation is probabilistic, with cytosine sometimes correctly incorporated (Shibutani et al. 1991; Cheng et al. 1992), and because other oxidative lesions block the polymerase (Dabney et al. 2013). Although a few early ancient DNA studies reported elevated *G* → *T* (Hofreiter et al. 2001; Briggs et al. 2007; Lamers et al. 2009), subsequent work concluded that cytosine deamination is the predominant source of postmortem miscoding lesions (Brotherton et al. 2007), and damage modeling has since centered on *C* → *T* and *G* → *A*, leaving oxidative damage comparatively understudied. However, it remains a recognized form of postmortem chemical damage in paleoproteomics (Warinner et al. 2022). We find evidence of oxidative damage in the Krestovka mammoth but not in the Antarctic subglacial precipitates, suggesting either that oxidative damage is common but its extent varies with environmental conditions, or that the subglacial samples carry too little damage to detect here given their much lower deamination than the Kap K samples. Because library preparation can also generate *G* → *T* (Costello et al. 2013), we confirmed the elevated *G* → *T* we report is not a lab preparation artifact (Supplementary Section S2.2). Without correction, this damage would bias downstream age estimates.

We developed a general approach to profile aDNA damage that produces nucleotide likelihoods to correct for damage within ratePlacer. With this correction, ratePlacer returned nonzero ages for Kap K, but the estimates did not agree across taxa: rather than the overlapping age ranges that concordance would produce, some ranges do not overlap, both overestimating and underestimating the published age (Table 4). We report each age as a range spanning the high and low *α* damage estimates and the divergence estimates used to calibrate the molecular clock.

**Table 4.** ratePlacer molecular age estimates and calibrated ages for *Betula* and additional most abundant genera from Kap København. The *Betula* consensus row is the chloroplast validation run against the Kjær et al. (2022) BEAST references; all other rows are nuclear read-based estimates using the damage-corrected approach, reporting both *α*_*hi*_ and *α*_*lo*_ estimates of the evolutionary-time parameter *α*.

| Taxon | Molecular age $\alpha_{hi}$ (95% CI) | Molecular age $\alpha_{lo}$ (95% CI) | Divergence (Ma) | 95% CI Converted age |
| --- | --- | --- | --- | --- |
| *Betula* (consensus) | 0.0004049 (0.0004047-0.0004050) | — | 61.1 | 1.631 to 1.632 Ma |
| *Betula* | 0.001504 (0.001498-0.001510) | 0.001451 (0.001444-0.001458) | 58-61 | $\sim 884$-963 ka |
| *Populus* | 0.009702 (0.009686-0.009718) | 0.01136 (0.01134-0.01137) | 35-52 | $\sim 2.44$-4.24 Ma |
| *Salix* | 0.01052 (0.01051-0.01054) | 0.01207 (0.01205-0.01208) | 35-52 | $\sim 2.95$-5.04 Ma |
| *Dryas* | 0.001977 (0.001968-0.001985) | 0.002027 (0.002019-0.002035) | 38-40.67 | $\sim 1.16$-1.29 Ma |

ratePlacer’s estimates are biased by two factors we can characterize but not eliminate: uncertainty in read placement and the treatment of damage. The short reads of aeDNA make the placement of each read within the reference tree uncertain (Bettisworth et al. 2025). The treatment of damage biases the estimate directionally. Leaving damage uncorrected pushes the age younger, because mismatches from damage inflate the apparent number of substitutions and lengthen ancient branches, whereas overcorrecting pushes it older, because mismatches that are in fact evolutionary substitutions are attributed to damage, shortening the aeDNA branches by making the reads more similar to the ancestral state. Our simulations modeled only deamination and so did not test ratePlacer under the oxidative damage we observed in the real data. ratePlacer also carries structural assumptions, each pointing to a needed extension. It assumes a strict molecular clock and ultrametric trees, so substitution rates cannot vary among lineages and calibrations from dated ancient genomes or remains cannot be incorporated. Relaxing these assumptions would allow both. It relies on a single reference topology, which cannot capture the distribution of genealogies produced by the coalescent. Finally, we did not benchmark the damage profiler against known damage probabilities, having confirmed only that its estimates track the observed positional substitution patterns. Such a benchmark remains needed.

Other limitations are shared with all molecular dating approaches rather than specific to ratePlacer. Recovered DNA is assumed to derive from a single time point, though processes that redistribute DNA within a deposit mean an unknown number of samples may in fact span multiple ages. A wide confidence interval can flag an age mixed sample, but the output remains a single average age, and resolving this is less a limitation specific to ratePlacer than an open challenge for the field, requiring methods that model multiple source ages, such as mixture models. The remaining shared biases arise from data processing and from the molecular clock, and act on the age in directions that are difficult to predict. Mapping discards reads that fail to align to the reference, disproportionately removing ancient endogenous reads with high mismatch counts, whether from true divergence, damage, or both, so the retained reads carry an artificially depleted substitution count that shortens branches and biases the age older. Taxonomic assignment then acts in two directions. The LCA step may fail to assign conserved reads to the order of interest, removing them, inflating the apparent substitutions per site, and biasing the age younger. Conversely, ngsLCA (Wang et al. 2022) retains reads only above a similarity threshold and, like mapping, removes reads with too many mismatches, so under high damage it biases the age older. We cannot determine which of these opposing effects dominates, it is likely sample dependent, and we recommend further work on the biases introduced by read mapping and assignment. Finally, the divergence estimates underlying the molecular clock affect the conversion from molecular age to years without a consistent direction. *Betula* and *Dryas* have relatively narrow reported divergence ranges, whereas *Salix* and *Populus* span wide ranges that widen our conversion accordingly. Because divergence estimates also differ across genetic compartments (nuclear, mitochondrial, chloroplast, or individual genes), we prioritized nuclear estimates with fossil calibrations. Acting together and in opposing directions, these processes may contribute to the cross taxa discordance reported above, though we cannot attribute it to any single one.

In practice, ratePlacer can estimate ages from as few as ~1,000 reads for older samples but closer to ~10,000 for younger ones, because older samples have typically diverged more from the reference and each read carries more of the substitutions that phylogenetic dating relies on. Correcting for damage is likely to raise these requirements, since the decomposition needs enough mismatches at each position to separate true substitution, damage, and sequencing error. We did not test these thresholds exhaustively, so they should be treated as rough guides rather than firm cutoffs. We recommend reporting ages across both the high and low *α* damage estimates to bound the effect of damage correction on the result.

More broadly, ratePlacer extends molecular dating to the short reads and mixed taxonomic composition of aeDNA, estimating ages that do not assume the recovered DNA is contemporaneous with the dated sediment. But damage, read placement, and data processing each bias these age estimates and must be accounted for. The putative oxidative damage we describe is one such bias: left uncorrected, it biases molecular evolutionary studies of aeDNA, ancient metagenomics, and low coverage aDNA, and it is easily overlooked where damage is subtle. Whether these modifications are driven by environment, age, or other processes remains open and will require direct experimental work rather than observation of aDNA alone. Together with the biases and assumptions detailed above, these remain open problems the field will need to work through as molecular dating of aeDNA develops. However, as long as these are kept in view, ratePlacer lays the groundwork to expand molecular dating to aeDNA. By combining many short reads across taxa without a consensus sequence, and by correcting for damage at low coverage, it dates aeDNA samples that earlier molecular dating methods could not. These estimates sharpen the sample ages that downstream ecological and evolutionary analyses rely on, such as tracking community turnover and species interactions over time.

## 4 Methods

### 4.1. ratePlacer

ratePlacer is a phylogeny-based method for estimating the molecular age of aDNA, developed specifically for ancient environmental DNA (aeDNA). ratePlacer takes advantage of pre-calculated fractional likelihoods to efficiently optimize the placement of reads along the assignment edge, similar to tronko (Pipes and Nielsen 2024). Reads are processed one at a time, computing a per-read likelihood *L*(*r*) for each read before combining these into a joint sample likelihood under the assumptions given below.

Reads from a sample are assumed to share a common age and to be conditionally independent given that age and the reference tree, allowing the joint sample likelihood to factor across reads. For a read assigned to edge *e* at a point that divides *e* into segments of length *t*_1_ and *t*_2_, and connected to that point by an edge of length *t*_3_, ratePlacer computes the phylogenetic likelihood

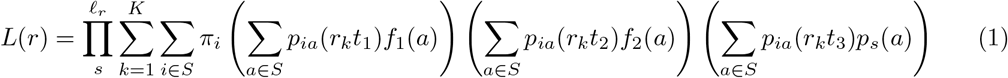

where *S* = {A, C, G, *T*} and i indexes the state at the read’s attachment node. The branch length *t*_1_ runs from the child node of edge *e* to the placement, *t*_2_ runs from the placement to the parent node of *e*, and *t*_1_ + *t*_2_ = *t*_*e*_ is held constant by the fixed reference tree. The branch length *t*_3_ runs from the ancient read leaf to its point of placement within the reference, and can be calculated during sample age estimation but is optimized over during single read age estimation. *f*_1_(*a*) and *f*_2_(*a*) are the fractional likelihoods at nucleotide *a* of the subtree descended from *e* and of the rest of the tree (with edge *e* and its descendant subtree removed), respectively (Fig. 5). *p*_*ij*_(*t*) is the GTR transition probability from *i* to *j* over branch length *t*, and *π*_*i*_ is the stationary frequency of nucleotide *i*. The index *k* runs over *K* discrete gamma rate categories with rate *r*_*k*_ (default *K* = 4) (Yang 1994), *s* indexes sites within the read of length *ℓ*_*r*_, and *p*_*s*_(*a*) is the probability of nucleotide *a* at site *s* of the read, derived from per-site damage profiles (see Section 4.3). Sites are assumed to evolve independently along the tree, allowing the per-read likelihood to factor across sites.

**Figure 4.**
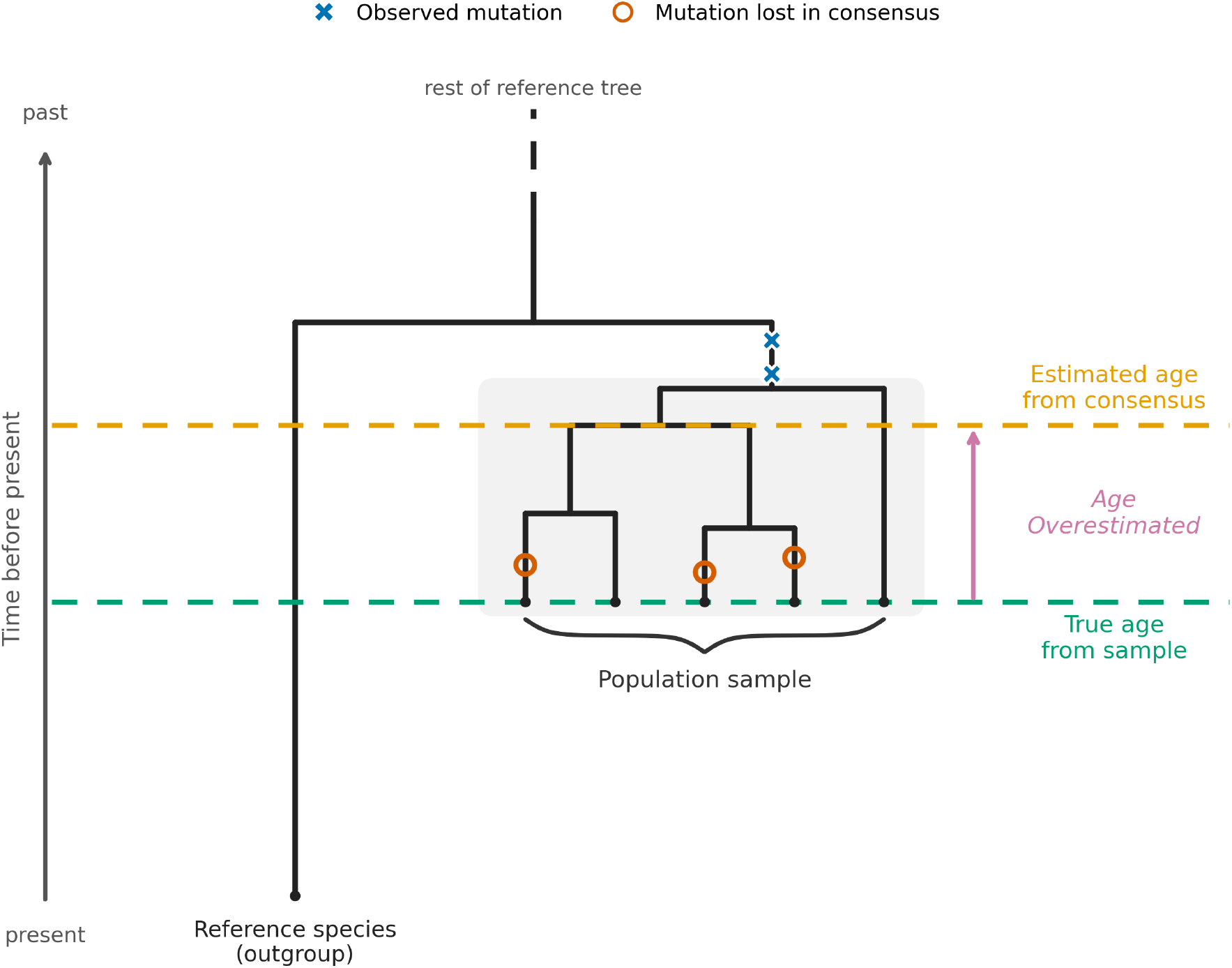
Building a consensus sequence from an ancient population biases age estimates older. Close-up of a reference phylogeny in which reads, though confidently assigned to a single species, are in fact sampled from a population. Reads are mapped to the closest reference species (outgroup), whose lineage survives to the present; the root connects to the rest of the reference tree (dashed). The sampled individuals form a coalescent whose tips mark the true age of the sample (green dashed line). Collapsing them into a single consensus sequence keeps only the mutations shared across the sample (observed, blue cross) and discards the recent diversity carried by individual lineages (lost, orange circle); because this pulls the sequence toward the ancestral state, the consensus is dated toward the coalescent root and the age is overestimated (yellow dashed line). We expect this to matter far more for young samples than for old ones: in a sample as old as Kap København, the lost diversity is small relative to the total branch length and should barely affect the estimated age, whereas in younger samples it makes up a larger fraction of the branch and can bias the estimate. The same effect arises when collapsing multiple species rather than a single population, where the greater discarded diversity makes the overestimate more extreme.

**Figure 5.**
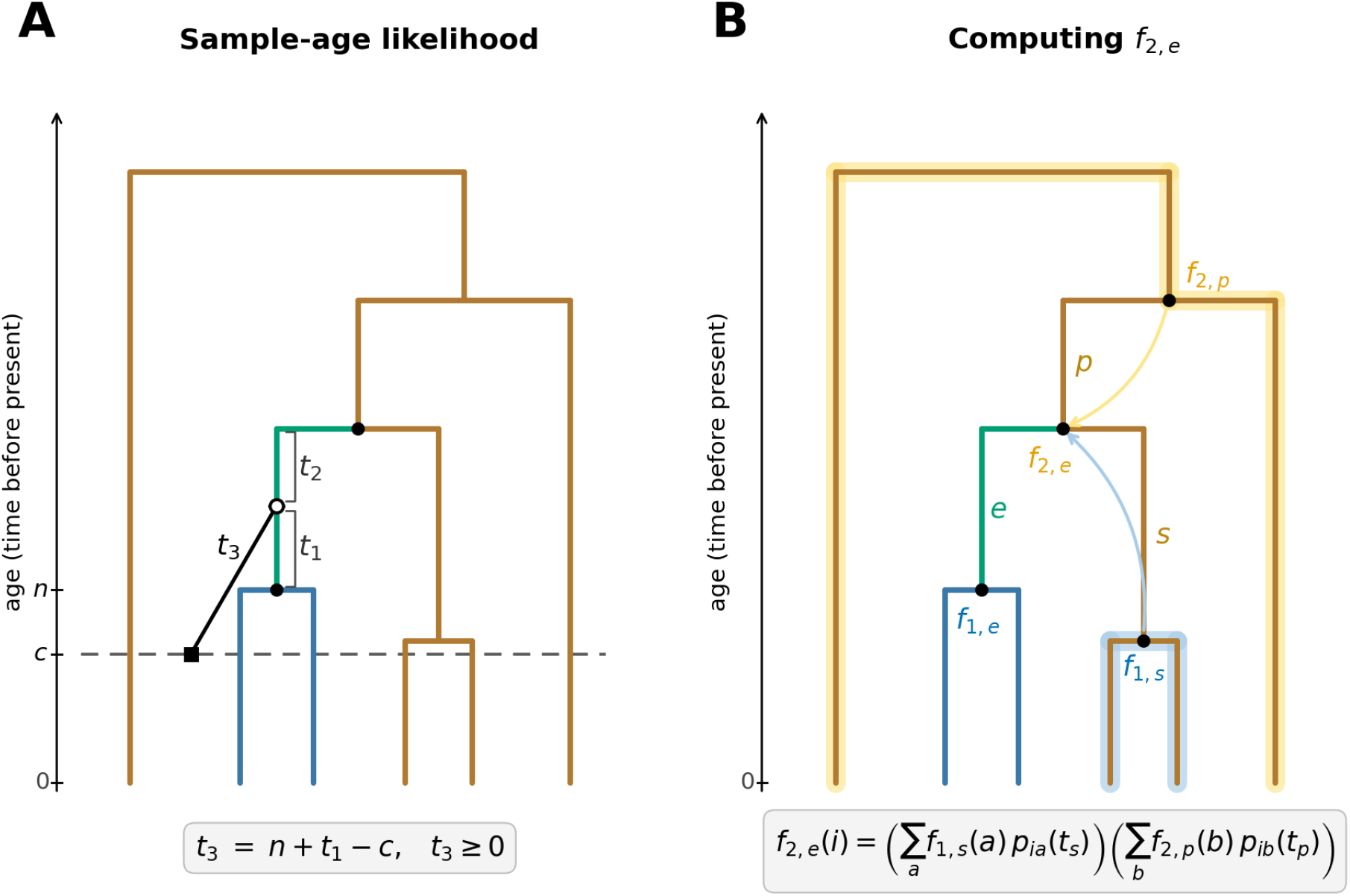
ratePlacer and gfl Schematic. A read (square node) is placed along an assignment edge *e* (green) of an ultrametric reference tree where *e* branches between a child node of age *n* to its parent node. The placement of the read in the tree divides *e* into segments *t*_1_ and *t*_2_ and the read has a branch length *t*_3_. The likelihood calculation is simplified by the pre-calculated subtrees with fractional likelihood *f*_1_ (in blue, subtree of the descendants of *e*) and *f*_2_ (in yellow, subtree of the rest of the tree) (calculated in gfl). (A) Sample age likelihood where sample age *c* (dashed line) is fixed and the read node lies on the age. *t*_3_ is now *n* + *t*_1_ − *a* (with *t*_3_ ≥ 0) and reduces the problem to a single free parameter *t*_1_ ∈ [max(0, *c* − *n*), *t*_*e*_] within the optimization of *c*. (B) Fractional likelihood calculation for *f*_2,*e*_, depicting how the recursion (light yellow highlight) and first postorder traversal (light blue highlight) are used to reduce the complexity.

Per-read placement is obtained by jointly optimizing the read placement *t*_1_ ∈ [0, *t*_*e*_] and the read branch length *t*_3_ ≥ 0 over the negative log-likelihood of Eq. 1 using Brent’s principal axis method (Brent 1973), a derivative-free algorithm for multidimensional minimization. When sample age is constrained, the placement optimization reduces to one dimension in *t*_1_ ∈ [max(0, *c* − *n*), *t*_*e*_], since the read branch length can be expressed as *t*_3_ = *n* + *t*_1_ − *c*, where *n* is the age of the child node of *e* and *c* is the constrained sample age (Fig. 5A). Sample-age estimation is implemented as two nested one-dimensional optimizations. The outer layer optimizes *c* using the sum of read log-likelihoods, and the inner layer optimizes *t*_1_ for each read with *t*_3_ constrained by *c*. Both one-dimensional layers use Brent’s method, a derivative-free algorithm for single dimension minimization. The primary output of ratePlacer is a maximum likelihood sample age estimate with a 95% confidence interval computed from the observed Fisher information at the maximum-likelihood estimate, evaluated via centered finite differences.

Fractional likelihoods are pre-calculated in a separate program, gfl (short for “get fractional likelihoods”), via a two-pass tree-traversal algorithm. Naively, computing *f*_1_ and *f*_2_ for every edge would require a separate tree traversal per edge, which is quadratic in the number of nodes. The twopass algorithm computes both quantities for all edges using a single postorder and a single preorder pass, which is linear in the number of nodes. A postorder traversal computes *f*_1_ for every edge using Felsenstein’s pruning algorithm (Felsenstein 1981). A preorder traversal then pulls fractional likelihoods down the tree to compute *f*_2_. *f*_2_ at edge *e* is calculated as

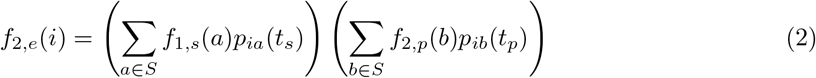

where *f*_1,*s*_(*a*) is *f*_1_ at nucleotide *a* of the sibling node, *f*_2,*p*_(*b*) is *f*_2_ at nucleotide *b* of the parent node, *t*_*s*_ is the sibling edge length, and *t*_*p*_ is the edge from the parent of *e* (Fig. 5B). The recursion is initialized by setting *f*_2_ at the root to 1 for all nucleotides. Input for gfl is an ultrametric reference and GTR+Γ parameters that are estimated with phangorn (Schliep 2011), with ultrametricity enforced via `minEdge(tree, enforce ultrametric = TRUE)` prior to fitting with `pml`.

ratePlacer and gfl are both implemented in C (Kernighan and Ritchie 2006) and available on GitHub (see Code Availability).

### 4.2. Analysis of DNA Damage Patterns

The damage analyses below operate on the BAM files and ancient/modern read classifications produced by the processing pipeline in Section 4.5.1.

Substitutions were counted directly from the curated BAM files by parsing the MD tag against the aligned reads using `pysam` (Heger et al. 2024). Counts were strand aware, correcting reverse strand mappings to the original forward orientation during analysis. For the ancient vs modern comparisons described below, counts were pooled across all read positions while position resolved counts were retained for the taxon-level analyses in Section 4.5.2. All counting and downstream Python and R analyses are available on GitHub (see Code Availability).

#### 4.2.1 Symmetric mutation flow

Under any time-reversible nucleotide substitution model, the detailed balance condition requires *π*_*i*_*Q*_*ij*_ = *π*_*j*_*Q*_*ji*_, where *π*_*i*_ is the stationary frequency of nucleotide *i* and *Q*_*ij*_ is the instantaneous substitution rate from *i* to *j*. Over an interval *t* and *n* aligned bases, the expected count of *i* → *j* substitutions is *nπ*_*i*_*Q*_*ij*_*t* and by detailed balance equals the expected count of *j* → *i*. We therefore expect observed counts to satisfy *N*_*ij*_ ≈ *N*_*ji*_ for all *i j* and *R*_*ij*_ = *N*_*ij*_*/N*_*ji*_ quantifies any asymmetry when it deviates from the expected ratio of 1. *R*_*ij*_ was computed for reads pooled by ancient/modern assignment and for the top ancient taxa.

To test whether asymmetries differed between ancient and modern reads, for each of the six substitution-type pairs we constructed a 2*×*2 contingency table with rows {ancient, modern} and columns {N_ij_, N_ji_} and applied a *χ*^2^ test of independence. Because read counts were large enough to produce significant *χ*^2^ results for negligible differences, we additionally report Cramér’s V as a measure of effect size. We use the relative magnitudes of Cramér’s V across substitution pairs to identify which asymmetries are most pronounced rather than to apply fixed effect-size thresholds.

#### 4.2.2 Observed-to-expected substitution ratios

In the absence of post-mortem damage and sequencing error, mismatches between reads and the reference should reflect only the evolutionary process, and the expected substituton counts can be calculated from the phylogenetic data using the estimated substitution model. With negligible sequencing error, then, in theory comparing observed mismatch counts to these expected counts should isolate any excess attributable to damage. For a given taxon, the expected count of *i* → *j* substitutions is *nπ*_*i*_*Q*_*ij*_*t*, where *n* is the number of aligned reference bases of type *i, π*_*i*_ the stationary frequency of nucleotide *i, Q*_*ij*_ the instantaneous substitution rate from *i* to *j*, and *t* the elapsed evolutionary time. Because *t* is unknown and acts as a global scaling factor that cancels in any comparison of ratios across substitution types, we set *t* = 1, treating the analysis as a relative comparison of substitution types within a taxon. The observed-to-expected ratio is then *O*_*ij*_ = *N*_*ij*_*/nπ*_*i*_*Q*_*ij*_. Assuming approximately equal elapsed time across ancient reads within a taxon, *O*_*ij*_ should be approximately uniform across substitution types in the absence of substitution-specific processes and departures from uniformity indicate post-mortem misincorporation. The reference base frequencies *π*_*i*_ and the aligned reference base count *n* were both computed from the reference bases at positions covered by the aligned reads, as recorded by `pysam` during substitution counting, ensuring the expected counts are computed over the same set of bases as the observed substitution counts. GTR+Γ parameters were estimated with `phangorn::pml` followed by `optim.pml` on the reference phylogeny (Section 4.1). Because GTR fitting requires a per-taxon multiple sequence alignment and phylogeny, these analyses were run independently on each phylogenetic group.

#### 4.2.3. Controls for taxonomic, bioinformatic, and sequencing bias

Two analyses were run to test whether observed asymmetries reflect post-mortem damage rather than sequencing error, non-reversibility of the substitution process, or taxonomic assignment.

##### Ancient vs. modern label permutation

To test whether the ratio *R*_*ij*_ differs between ancient and modern reads, we tested the null hypothesis that *R*_*ij*_ has the same distribution in ancient and modern reads. Ancient and modern labels for 192,258,396 ancient and 5,404 modern reads from the Kap København samples were randomly permuted 10,000 times. For each permutation and each substitution pair, *R*_*ij*_ was computed in the permuted ancient and modern pools, and the test statistic *t*_*ij*_ = *R*_*ij*_(ancient) − *R*_*ij*_(modern) was recorded, yielding a null distribution per pair. Empirical twosided p-values were computed as *p* = (1+#{|*t*_null_| ≥ |*t*_obs_|})/(1+10,000). Rejecting this null indicates that post-mortem processes contribute to the asymmetry, whereas failure to reject indicates that sequencing error, non-reversibility of the substitution process, or the taxonomic composition of the two pools could account for the observed pattern.

##### Watson-Crick pair test

The damages of interest here arise as strand-specific DNA lesions from damage processes such as cytosine deamination and guanine oxidation. These modifications are converted into apparent substitutions during library construction. In double-stranded libraries, the strand of origin can be lost during sequencing and a true *i* → *j* substitution can be observed as its complement on the opposite strand. Under unbiased sequencing, counts of complement substitution pairs should occur at a 1:1 ratio and a deviation from 1 indicates strand specific sequencing or alignment bias. We computed ratios on samples from Kap København, which only contain double-stranded libraries. Single-stranded libraries were excluded from this analysis as the substitution strand of origin for damage is preserved and so complement ratios are not expected to be 1:1, even under unbiased sequencing.

### 4.3. DNA Damage Nucleotide Error Model

#### 4.3.1. Generative Model

Mismatches between the reference and observed reads result from three independent processes: (i) evolutionary substitutions accumulated since divergence from the reference, (ii) a position specific post-mortem or chemical DNA damage, and (iii) base-calling errors introduced in sequencing. Of these processes, damage is position specific in the read, while we assume sequencing error and evolutionary processes are not. To estimate the damage component, we model each process as a 4*×*4 row-stochastic transition probability matrix in the order {A, C, G, T}. The damage matrix is the primary object of inference.

Each read is collapsed into *K* bins. By default *K* = 21 with one bin per base for the first and last 10 positions, plus a single aggregated bin for the interior. This reflects that the 5’ and 3’ termini experience variation in damage while the middle section remains relatively stable. For each *k* ∈ {1, …, *K*}, the observed mismatch matrix 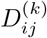 is the empirical proportion of reads in bin *k* where *j* is the observed nucleotide and *i* is the nucleotide in the reference sequence. Let *A*(*α*) represent the evolutionary substitution matrix shared across all bins, where *A*(*α*) = *e*^*αQ*^. *Q* is the instantaneous rate matrix estimated from fitting a phylogenetic model on the modern reference sequences closely related to the taxon of interest under a GTR+Γ model of evolution. *α, α* ≥ 0 is an unknown scalar that represents the elapsed evolutionary time between the modern references and ancient reads. *α* is also upper-bounded at *α*_*max*_ = 1*/max*_*i*_(−*Q*_*ii*_) to ensure a valid transition matrix. Let 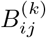 be the probability that post-substitution nucleotide *i* is damaged to nucleotide *j* for bin *k*. Let *C* represent the sequencing error rates estimated from the average base quality scores among all reads and positions. For nucleotide *i*, the mean error probability *e*_*i*_ is calculated from the Phred scores of all the reads with observed base *i*, giving *C*_*ii*_ = 1 − *e*_*i*_, and *C*_*ij*_ = *e*_*i*_*/*3 for *i ?*= *j*, assuming errors are equally likely among the three alternative bases.

These three processes act sequentially and independently on each site, giving the generative model in matrix form as

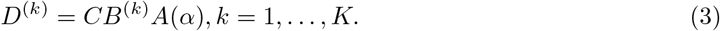

#### 4.3.2. Estimation Procedure

We estimate damage rates using maximum likelihood under Equation 3, where all matrices are row-stochastic and entries of *B*^(*k*)^ are constrained to be non-negative. The likelihood is multinomial and for a fixed *α*, the log likelihood for bin *k* is

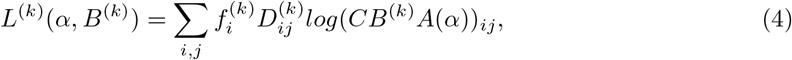

where 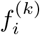 is the background nucleotide frequency of *i* for bin *k*. For fixed *α*, each *L*^(*k*)^(*α, B*^(*k*)^) depends only on data from bin *k*, so the *K* inner problems for *B*^(*k*)^ decouple and are solved independently via sequential least squares programming (SLSQP). The bins are coupled through the shared *α*, which is estimated jointly by maximizing the profile log-likelihood 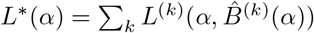.

Since *L*^∗^(*α*) is flat over the joint feasible interval *F, α* is not identifiable within *F*, and it is not possible to find a unique maximum likelihood estimate for {B^(k)^} from data in *F* alone. Therefore the feasible interval for bin *k* is defined as

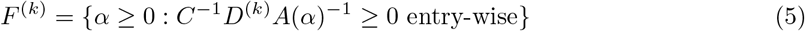

and defines the range of *α* that an analytical solution for *B*^(*k*)^(*α*) = *C*^−1^*D*^(*k*)^*A*(*α*)^−1^ can be found such that *B* is non-negative. Since *α* is shared across all categories, the joint feasible set is

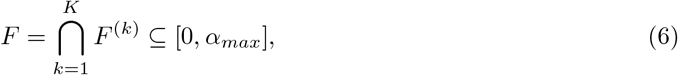

with *α*_*max*_ as defined in Section 4.3.1. Within this interval, all *α* within *F* have equally supported damage estimates. Outside the bounds of *F*, the constrained solver returns valid {B^(k)^} with some entries as 0 and returns a strictly worse likelihood. Using Brent’s root-finding algorithm applied to 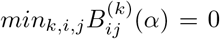, the bounds *α*_*lo*_ and *α*_*hi*_ of *F* are located. 95% confidence intervals are then constructed via the likelihood ratio criterion, finding the values of *α* where 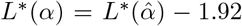 (corresponding to the 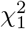 threshold) using Brent’s root-finding algorithm. Minimum and maximum damage estimates *B*^(*k*)^(*α*_*loCI*_) and *B*^(*k*)^(*α*_*hiCI*_) are evaluated at the CI bounds and bracket the range of damage profiles consistent with the data. *B*^(*k*)^(*α*_*loCI*_) can be solved analytically using *B*^(*k*)^ = *C*^−1^*D*^(*k*)^*A*(*α*)^−1^ and *B*^(*k*)^(*α*_*hiCI*_) is found using the multinomial log-likelihood solver with SLSQP.

#### 4.3.3. Nucleotide Likelihoods from Damage Matrix

Having estimated the damage profiles 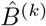, we can account for post-mortem damage when computing nucleotide likelihoods for input to ratePlacer’s molecular clock dating. Molecular age estimation depends on resolving evolutionary substitutions from the observed bases. Without correction, damage-induced mismatches would be conflated with true substitutions and bias the estimates. By combining the damage estimates with per-base sequencing error, we compute likelihoods over the true pre-damage base state at each position, isolating the evolutionary signal used for dating.

For each observed base in the dataset, we compute a nucleotide likelihood over the true unobserved base *Y*_*k*_ ∈ {A, C, G, *T*}, defined as the state after evolutionary substitution but prior to damage and sequencing error. Let *X*_*k*_ ∈ {A, C, G, *T*} be the observed base at a read position assigned to bin *k*, and let *e*_*k*_ be its per-base Phred-derived error probability. Unlike the averaged error matrix *C* used during estimation, here we use the base-specific error probability directly, constructing a per-observation error 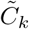 with 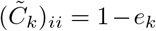 and 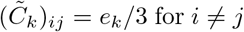 for *i* ≠ *j*, assuming sequencing errors are uniformly distributed across the three alternative bases.

The nucleotide likelihood is then

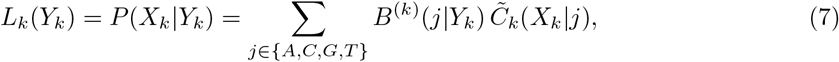

where *B*^(*k*)^ is evaluated at either *α*_*loCI*_ or *α*_*hiCI*_. As the two CI bounds bracket the range of damage profiles consistent with the data, computing likelihoods under both propagates uncertainty in the damage estimate through to ratePlacer, yielding a corresponding range of age estimation. These likelihoods are computed for every base in the dataset and supply *p*_*s*_(*a*) in Eq. 1 when ratePlacer is run on damage-corrected data.

### 4.4 Simulation Pipeline

To test how well ratePlacer performs with aeDNA data, we simulated reads across a range of conditions: read length, read count, number of taxa, data filtering, and damage level. Three ages were tested (10^5^, 10^6^, and 10^7^ years), converted from branch lengths under an assumed rate of 10^−9^ subs/site/year for interpretability. ratePlacer operates and reports on branch lengths. Unless otherwise noted, each condition was simulated 1,000 times.

A single phylogeny containing twenty-two taxa was used as the reference across all simulations. In the single-source case, reads were sampled from one ancient leaf node. In the mixed case, reads were sampled from three ancient nodes, each placed on a different branch within a single clade of the reference tree, so the three sources shared an age but differed by the lineages within the tree. Starting from this phylogeny with ancient sequences set to the target age within the reference, full length sequences were simulated using IQ-TREE Alisim (Ly-Trong et al. 2022) under a GTR+Γ model. An Rscript was used to convert the reference tree and parameters into a ratePlacer readable format.

Reads were sampled from the simulated ancient sequence at the specified length and count. In the no-damage condition, reads were sampled directly from the ancient sequence without sequencing error or substitution damage. When damage was specified, substitution damage and sequencing error were introduced using gargammel (Renaud et al. 2017), which combines deamSim for damage and ART (Huang et al. 2012) for sequencing error. With deamSim, default parameters were used except `-d 0.001`, the rate of deamination on double-stranded regions. With ART, we used `art illumina` with `-ss HS25 -amp -c 1 -l L` to apply sequencing error without modifying the read lengths, default parameters otherwise. When damage correction was applied, damage rates were estimated per-replicate using the method described in Section 4.3. ratePlacer’s internal branch assignment requires a starting node in the reference tree. Read assignment was performed using either `tronko-assign` or by supplying a node index in [1, 2 *× n*_*references*_] directly and using ratePlacer’s maximum likelihood algorithm to bypass tronko entirely. When `tronko-assign` was used, `tronko-build` was first applied to the reference sequences and ultrametric tree. ratePlacer was run in sample age estimation mode across all conditions. For multi-taxa cases, reads from all taxa were pooled together. Per-branch merging of reads was applied with a minimum cutoff of 500bp for inclusion in age estimation. Results were aggregated across replicates and visualized as histograms using ggplot2 (Wickham 2016) in R (R Core Team 2021).

For a subset of conditions (damage, low coverage, and multiple taxa) we compared ratePlacer estimates to BEAST X (Baele et al. 2025) tip dating. A general BEAUti template was generated and modified by a custom Python (Van Rossum and Drake 2009) script to substitute sequences for each replicate. Following Kjær et al. (2022), simulated aeDNA reads were merged into a single consensus sequence. For multi-taxa cases, per-taxon consensus sequences were also generated to enable comparison against the pooled single-sequence case. Reference sequences were trimmed to match the consensus alignment. Tip-dated sequences were assigned a uniform prior on age. The reference taxon set was assigned tip-date ages of 0. A normal prior on root age was set to 26.46 with a standard deviation 0.5. A strict molecular clock was assumed under GTR+Γ with all parameters estimated. A constant-size coalescent tree prior was used with a 1/x prior. One MCMC chain of 100,000,000 iterations logged every 1,000 iterations was run per replicate with a 10% burn-in. Convergence was verified on a subset of runs using Tracer (Rambaut et al. 2018). Point estimates were taken as the median of the ancient tip age. Due to computational cost of BEAST X, this comparison used 100 replicates rather than 1,000.

### 4.5. Ancient DNA Analysis

Three datasets were analyzed in this paper: ancient environmental DNA (aeDNA) from Kap København (Kjær et al. 2022), aeDNA from Antarctic subglacial precipitates (De Sanctis et al. 2025a), and aDNA from a Krestovka mammoth molar (approximately 1.65 million years old) (Valk et al. 2021). Not all datasets were used in every analysis. Kap København was processed from raw reads and used in all downstream analyses, the Krestovka mammoth was reprocessed from raw reads for single-taxon damage analysis only, and the Antarctic subglacial samples were obtained as processed BAM and LCA files for the aggregate symmetry analysis only. All custom analysis scripts for this paper are available on GitHub (see Code Availability).

#### 4.5.1. Processing of Ancient DNA

##### Kap København

For analyses in this study, we considered only the 29 Kap København libraries which were shotgun sequenced (as opposed to capture) and which were used in the main figures of Kjær et al. (2022), as well as the negative controls from the original study.

Raw reads from (Kjær et al. 2022) were downloaded from ENA project PRJEB55522. Quality control and preprocessing were performed using fastp (Chen et al. 2018) with the following parameters: paired-end reads were merged using the `-m` flag, with custom double-stranded adapter sequences specified via `--adapter fasta`. Deduplication was enabled (`-D`) with accuracy set to 5 (`--dup calc accuracy 5`). PolyG and polyX tail trimming were forced (`-g -x`) with minimum lengths of 5 bases (`--poly g min len 5 --poly x min len 5`). We retained only bases with scores ≥ 30 (`-q 30`) and reads with average quality ≥ 25 (`-e 25`), with a minimum read length threshold of 30 bp (`-l 30`). Additional filters included low complexity filtering (`-y`), base correction in overlapped regions (`-c`), and overrepresented sequence analysis (`-p`). Further quality control and deduplication of exact match and exact substring reads was performed using SGA with dust filtering (threshold=30), minimum read length of 30 bp, and kmer-based filtering disabled (Simpson and Durbin 2012). Final quality assessment was conducted using FastQC (Andrews 2010).

We mapped against two reference databases composed of eukaryotic and microbial reads (Supplementary Section S2.3.1). Reads were mapped in parallel with Bowtie 2 `-k 100` and default parameters otherwise (Langmead and Salzberg 2012). Bams were merged for each of the two databases separately, query-sorted, and unnecessary headers were removed with bamsort (Durbin 2025). Reads aligning to any references in GTDB were removed from the eukaryotic bams. This yielded a microbial and a eukaryotic bam for each library.

Next, for each library, we ran ngsLCA (Wang et al. 2022) with `-fix-ncbi 0, -simscorelow 0.95, -simscorehigh 1.0` and default parameters otherwise, for each of the microbial and eukaryotic bam files separately. Next we ran bamdam (De Sanctis et al. 2025b) `shrink` and `compute` with default parameters, with `--stranded ds` (double stranded libraries) and with `--upto phylum` for the microbial bams.

Using a custom script, taxa were classified as ancient if they had mean damage greater than 0.4, DUST less than 7, and more than 10,000 reads, and as modern if they had mean damage less than 0.1, DUST less than 7, and more than 500 reads. Modern taxa with mean read counts higher in control samples were removed as contaminants (Supplementary Section S2.2). Reads and alignments for retained taxa were extracted with `bamdam extract --only-top-alignment` and default parameters otherwise.

For single-taxon analyses, retained reads were remapped with Bowtie 2 (same parameters as above) to a per-genus database containing all available nuclear, mitochondrial, and chloroplast genomes for the focal genus and its closest outgroup (Supplementary Section S2.4). Reads were assigned to a specific genetic compartment only if they did not multimap across compartments, to the outgroup, or to multiple positions within a single genome.

##### Krestovka mammoth molar

Raw reads were downloaded from ENA (accession PRJEB42269) and trimmed, merged, and deduplicated with fastp and sga following the same pipeline as Kap København. Reads were mapped with Bowtie 2 against a database of elephant and human nuclear and mitochondrial reference genomes (Supplementary Section S2.3.2) and assigned to genetic compartments using the same multimapping criteria as the Kap København single-taxon analyses.

##### Antarctic subglacial precipitates

Processed BAM and LCA files, along with ancient and modern taxonomic assignments, were obtained from the authors. Reads from subglacial precipitates for downstream analyses were extracted with `bamdam extract --only-top-alignment` and default parameters otherwise (Supplementary Section S2.5).

#### 4.5.2. Supplementary Damage Analysis

Single-taxon damage analyses were applied to the nuclear reads of the top genus-level hits from Kap København. GTR+Γ parameters were estimated from alignments of the nuclear reference genomes trimmed to positions overlapping the ancient reads from Kap København. Aggregate and positional substitution counts and observed-to-expected ratios were computed using `calc_subs.py`.

The full aggregate symmetry analysis between modern and ancient reads (Section 4.2.1) along with bias controls (Section 4.2.3) was applied to Kap København. Antarctic subglacial samples were included as an additional visual comparison of ancient and modern substitution patterns without statistical testing, along with nuclear reads assigned to Krestovka mammoth.

#### 4.5.3. Molecular Dating of Kap København

ratePlacer was applied to one chloroplast genus (*Betula*) and four nuclear genera (*Betula, Salix, Populus*, and *Dryas*). The chloroplast analysis provides a direct comparison to the BEAST tip-dating estimate of Kjær et al. (2022) and uses their input sequences directly. The nuclear analyses use reads processed in Section 4.5.1.

In all cases, reference trees were estimated with RAxML under GTRGAMMA model (version 8.2.12) and made ultrametric with GTR+Γ parameters estimated using phangorn. Fractional likelihoods for the reference phylogenies were precomputed with gfl (Section 4.1).

##### *Betula* chloroplast

Reference sequences (thirty *Betula* and one *Alnus* as outgroup; Supplementary Section S2.3.3) and the consensus chloroplast sequence representing the Kap København ancient sequence were extracted from the BEAUti XML provided by Kjær et al. (2022).

References were used to build the ratePlacer reference database as described above. Because the ancient sequence is a consensus sequence from sites with ≥ 20× depth as defined by Kjær et al. (2022), no damage error profile was generated. ratePlacer was run on the consensus with default parameters.

##### Nuclear pipeline

For each genus, reference genomes were aligned with progressive Cactus (version 9.1.2) (Armstrong et al. 2020) using default parameters. *Populus* and *Salix* share a single Cactus alignment containing both genera but were processed as separate ratePlacer runs. *Betula* and *Dryas* each have their own Cactus alignment. The HAL output was converted to MAF with `hal2maf` (Hickey et al. 2013) using a backbone reference from the focal genus (Supplementary Section S2.3.3), as mapping rates varied with reference choice. The MAF was stitched together into a single multiple sequence alignment with a custom script. Reads classified as nuclear (Section 4.5.1) were remapped to the aligned references, and the references were trimmed to read-overlapping positions. Per-site nucleotide likelihoods incorporating damage profiles were generated with a custom Python script following the procedure described in Section 4.3. ratePlacer was then run with default parameters on both hi and lo *α* damage estimates.

##### Age estimates without the damage model

Prior to developing the damage model (Section 4.3), we evaluated three dating approaches in which per-site nucleotide likelihoods were derived from base quality scores alone. These differed in how post-mortem deamination was handled when reads were compared to the reference: (i) masking potential deamination sites at the first and last seven bases of each read, (ii) masking all transitions, and (iii) trimming the first and last seven bases of each read. ratePlacer was run on each configuration with default parameters. These approaches did not adequately account for damage and produced biased age estimates (Section 2.3), motivating the damage model used in the main pipeline.

## Acknowledgments

We would like to thank John Huelsenbeck, John Lo, and Ruairidh Macleod for their helpful conversations and guidance.

## Author Contributions

MLK and RN developed ratePlacer. MLK performed the molecular dating of Kap København, simulations, symmetry analysis, nucleotide likelihood model, and wrote the paper, with input from all other authors. LP updated tronko to be compatible with ratePlacer. BDS ran the Kap København remapping and provided guidance on a(e)DNA. RN developed the damage profile method and provided supervision on all of analyses.

## Code Availability

Source code for ratePlacer and gfl, along with all simulations, pipeline, and analyses from the manuscript are available at the following GitHub repositories.

The source code for ratePlacer, gfl, simulations, and helper scripts can be found at: https://github.com/lemmonquiche/ratePlacer

The code for the symmetry analysis can be found at: https://github.com/lemmonquiche/ancientDNASymmetry

## AI Use Statements

During the preparation of this work, Claude 4.6-8 was used to assist with editing the manuscript for clarity and parts of the code/data visualization scripts; the authors reviewed and edited all the content for accuracy and take responsibility for the accuracy and integrity of the work.

