## Supplemental Information for "Molecular Clock Dating of Ancient Environmental DNA Reveals Damage Beyond Deamination"

### S1 Supplementary Figures

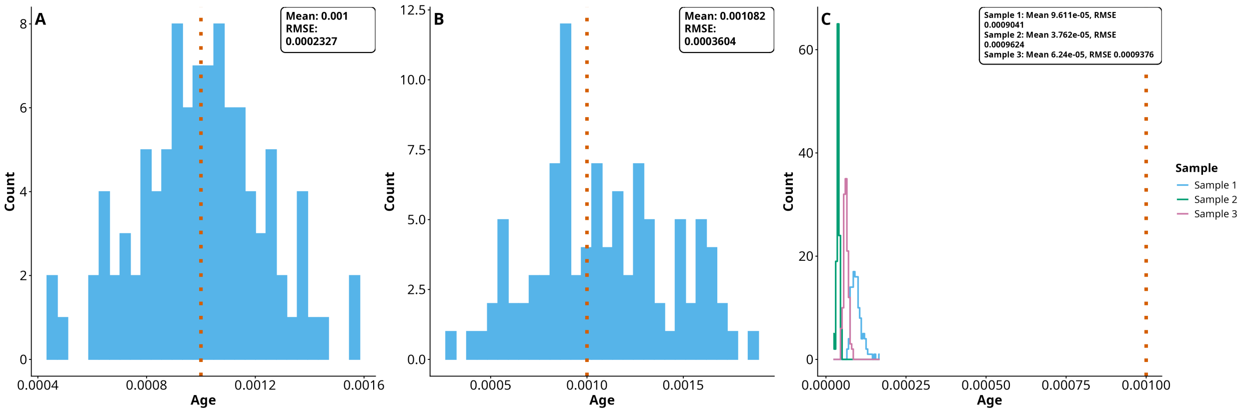

**Figure S1: Distributions of 100 BEAST X tip-dating age estimates.** The orange dashed line represents the true age, 0.001 substitutions/site in each case, with per-panel mean and RMSE are inset. A) Age estimates of the full 125kbp ancient sequences. B) Consensus sequence age estimate of the downsampled ancient sequence with no damage. C) Age estimates of the mixed taxonomy sample with damage, separating the reads and building a consensus sequence per ancient taxon.

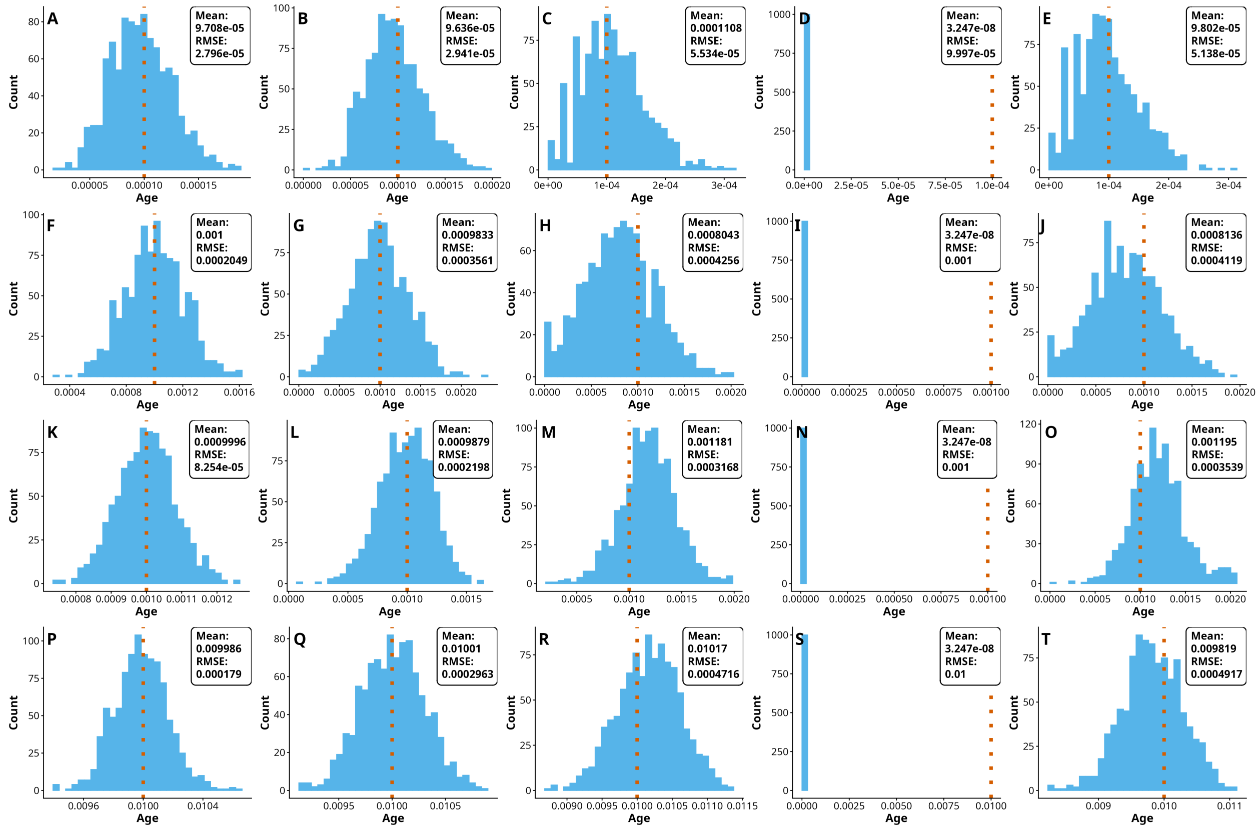

**Figure S2: Distributions of 1,000 ratePlacer age estimates across age, damage, number of taxa, and coverage.** The orange dashed line represents the true age with per-panel mean and RMSE are inset. (A-E) age 0.0001, (F-O) age 0.001, and (P-T) age 0.01. (K-O) samples have three taxa, while the rest are single taxon. (A, F, K, P) are age estimates on the full 125kbp ancient sequences, the rest are downsampled to 1,000 reads of 50bp length. (B, G, L, Q) True placement and no damage. (C, H, M, R) Tronko placement and damage. (D, I, N, S) Tronko placement with uncorrected damages. (E, J, O, T) Tronko placement with damage correction.

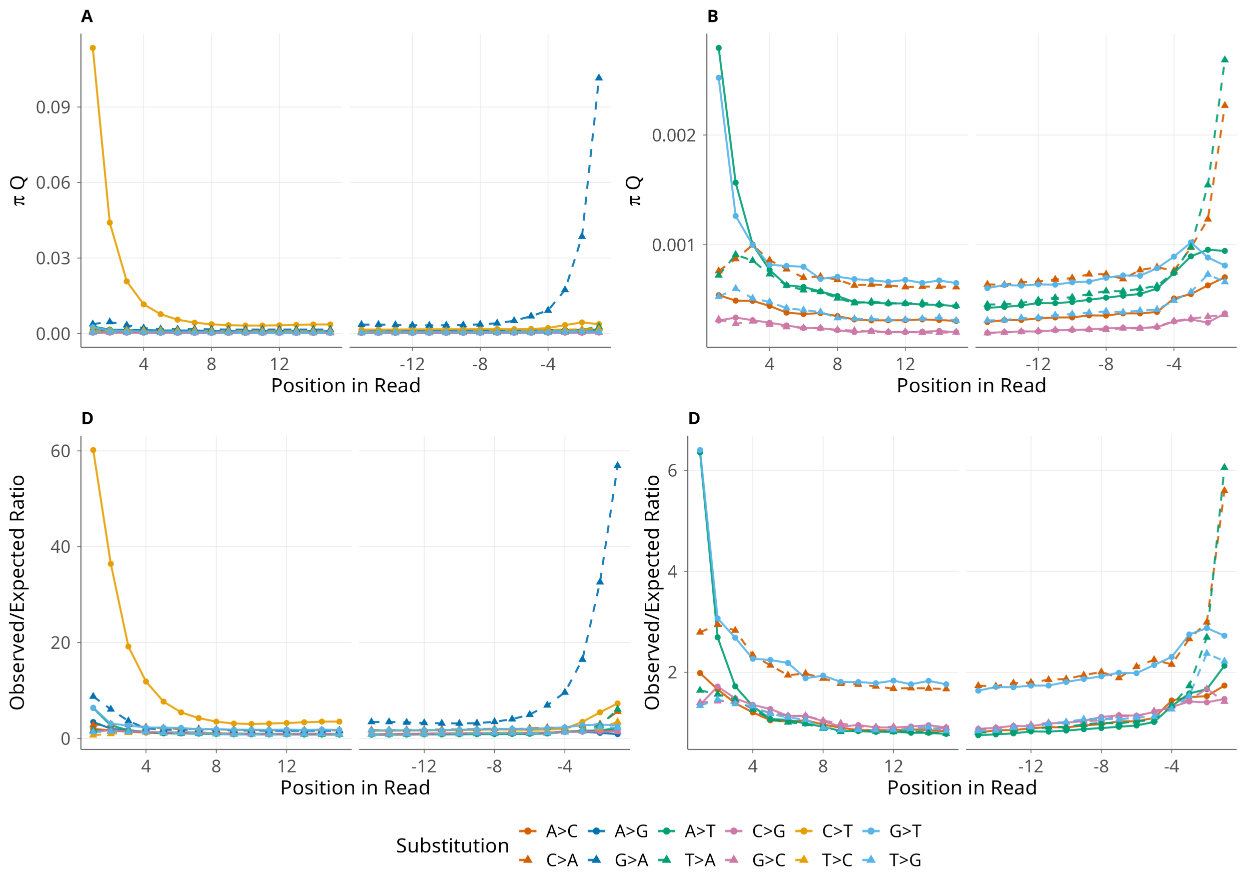

**Figure S3: Positional substitution patterns for *Dryas* reads at the first and last fifteen positions.** Panels (A) and (C) show all substitutions; panels (B) and (D) show non-deamination damage substitutions ($C \to T$ and $G \to A$ are removed). Deamination damage substitutions are removed from panels (B) and (D) due to differences in scale. Panels (A,B) show observed rates ($\pi Q$); panels (C,D) show observed-to-expected ratios, with the same scaling used for *Dryas* in Table 1. Solid lines denote one direction of each substitution pair and dashed lines denote the reverse (e.g., solid $C \to T$ vs. dashed $T \to C$). Under the symmetry assumption, matched dashed and solid lines of the same color should overlap in (A,B), and all lines should overlap in (C,D); gaps between matched pairs indicate violations of symmetry. Elevation above the background for both deamination ($C \to T$, $G \to A$) and oxidative damage ($G \to T$, $C \to A$) substitutions is visible across the interior of the read in addition to the terminal positions.

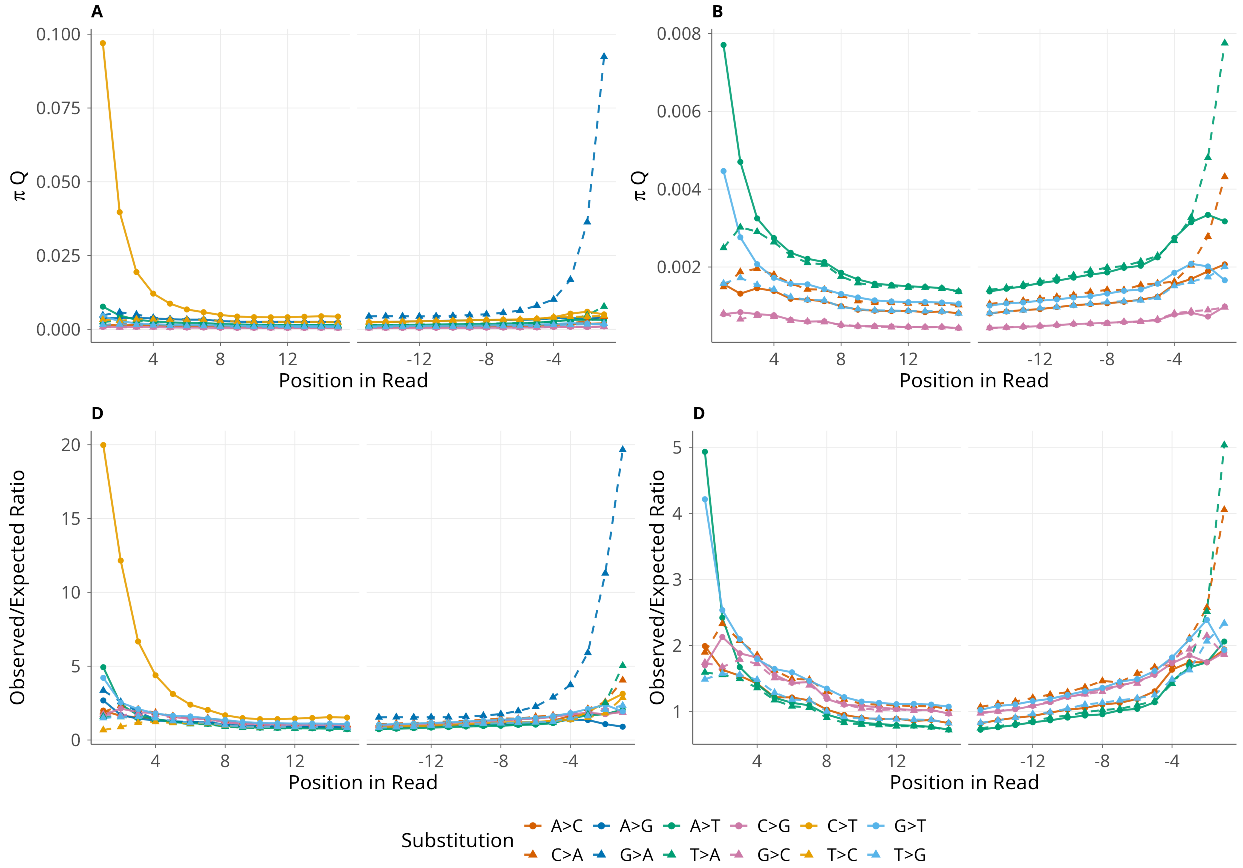

**Figure S4: Positional substitution patterns for *Salix* reads at the first and last fifteen positions.** Panels (A) and (C) show all substitutions; panels (B) and (D) show non-deamination damage substitutions ($C \to T$ and $G \to A$ are removed). Deamination damage substitutions are removed from panels (B) and (D) due to differences in scale. Panels (A,B) show observed rates ($\pi Q$); panels (C,D) show observed-to-expected ratios, with the same scaling used for *Salix* in Table 1. Solid lines denote one direction of each substitution pair and dashed lines denote the reverse (e.g., solid $C \to T$ vs. dashed $T \to C$). Under the symmetry assumption, matched dashed and solid lines of the same color should overlap in (A,B), and all lines should overlap in (C,D); gaps between matched pairs indicate violations of symmetry. Elevation above the background for both deamination ($C \to T$, $G \to A$) and oxidative damage ($G \to T$, $C \to A$) substitutions is visible across the interior of the read in addition to the terminal positions.

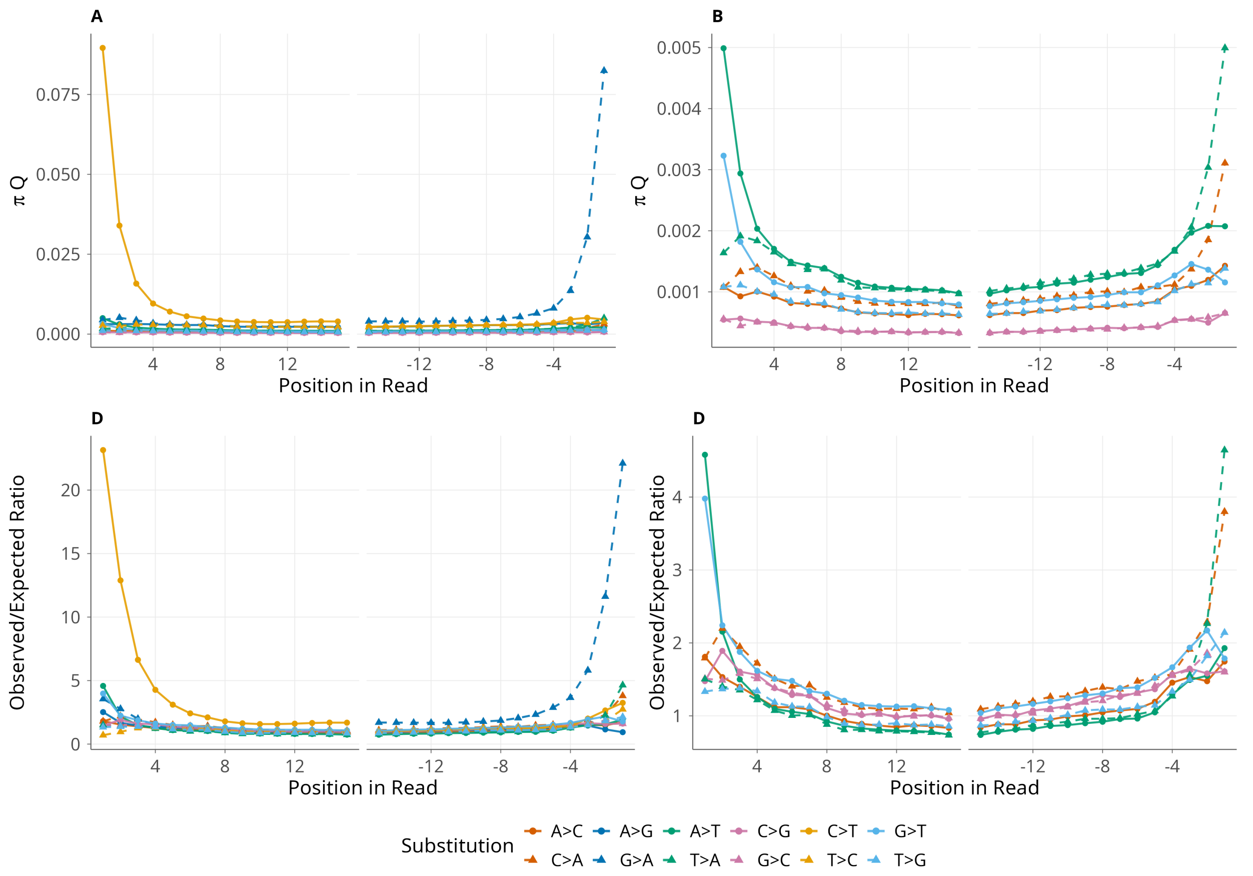

**Figure S5: Positional substitution patterns for *Populus* reads at the first and last fifteen positions.** Panels (A) and (C) show all substitutions; panels (B) and (D) show non-deamination damage substitutions ($C \to T$ and $G \to A$ are removed). Deamination damage substitutions are removed from panels (B) and (D) due to differences in scale. Panels (A,B) show observed rates ($\pi Q$); panels (C,D) show observed-to-expected ratios, with the same scaling used for *Populus* in Table 1. Solid lines denote one direction of each substitution pair and dashed lines denote the reverse (e.g., solid $C \to T$ vs. dashed $T \to C$). Under the symmetry assumption, matched dashed and solid lines of the same color should overlap in (A,B), and all lines should overlap in (C,D); gaps between matched pairs indicate violations of symmetry. Elevation above the background for both deamination ($C \to T$, $G \to A$) and oxidative damage ($G \to T$, $C \to A$) substitutions is visible across the interior of the read in addition to the terminal positions.

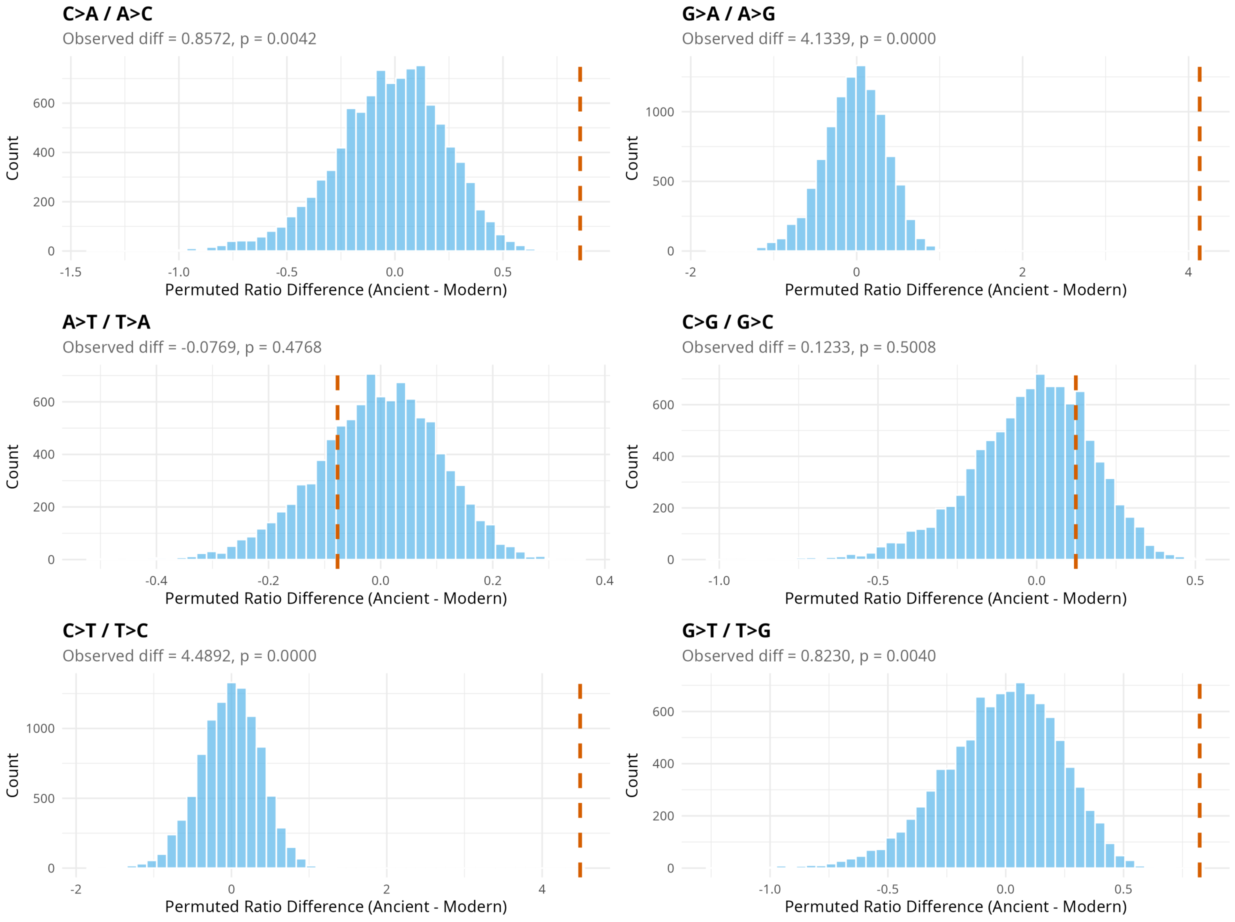

**Figure S6: Permutation test of taxonomic-label influence on substitution ratio differences.** For each symmetric substitution pair, ancient and modern labels were randomly reassigned and the difference in mean substitution ratio (ancient - modern) recomputed over 100,000 iterations; the histogram is the resulting permutation null distribution and the dashed red line marks the observed difference. Per-panel annotations give the observed difference and permutation p-value. A red line far in the tail of the null indicates the observed elevation is not explained by taxonomic composition.

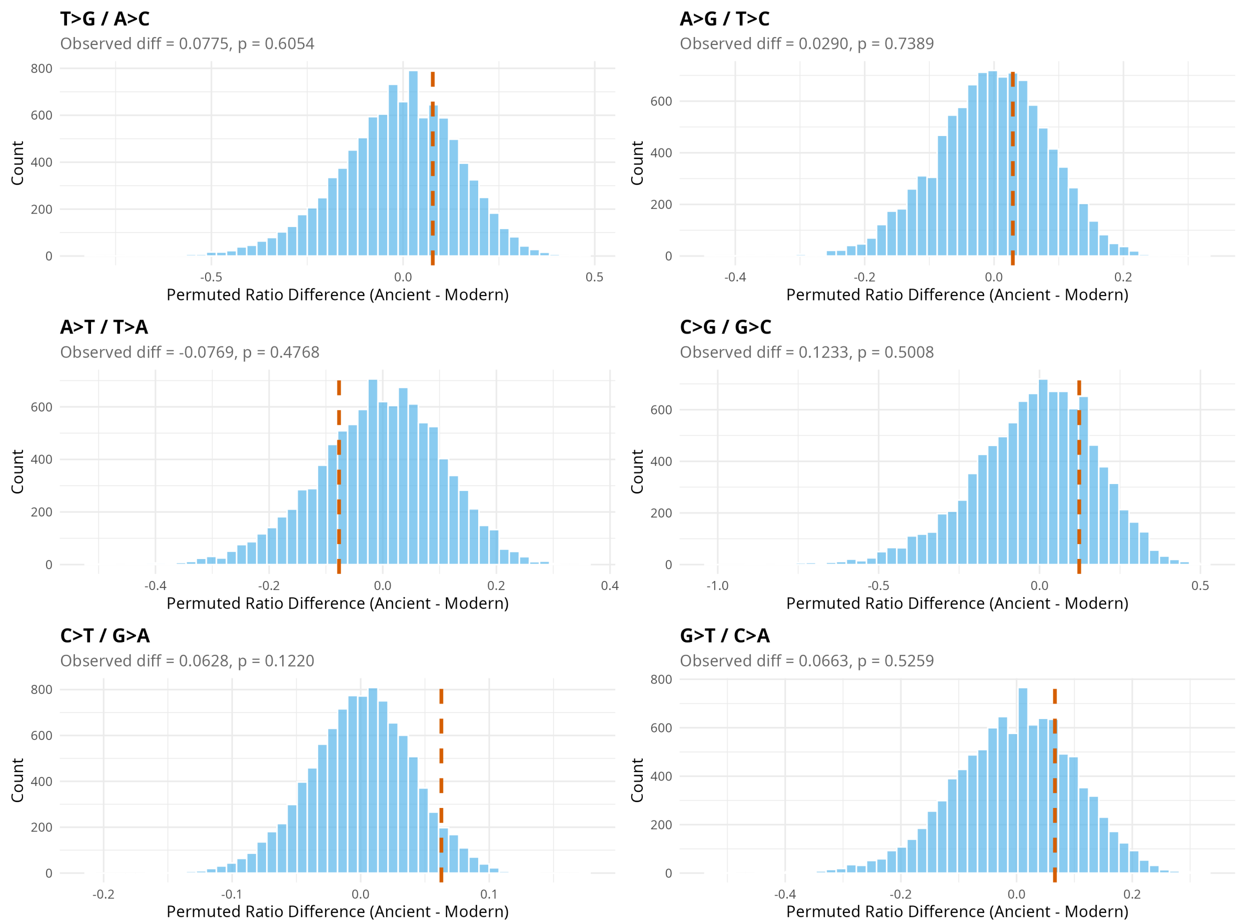

**Figure S7: Permutation test of taxonomic-label influence on substitution ratio differences.** For each Watson-Crick substitution pair, ancient and modern labels were randomly reassigned and the difference in mean substitution ratio (ancient - modern) recomputed over 100,000 iterations; the histogram is the resulting permutation null distribution and the dashed red line marks the observed difference. Per-panel annotations give the observed difference and permutation p-value. A red line far in the tail of the null indicates the observed elevation is not explained by taxonomic composition.

### S2 Supplementary Information

#### S2.1 Non-Time Reversible Parameters in Observed-to-Expected Ratios using UNREST Models

We tested whether our observed-to-expected ratios calculated by GTR parameters held under a non-reversible substitution model, since substitution biases can be better explained by such models (Sianga-Mete et al. 2025). We used IQ-TREE (Minh et al. 2020) to estimate UNREST (Yang 1994) model parameters, which we used to calculate expected counts for the observed-to-expected ratios (Table S1). We found that overall, model choice does not explain the elevated $G \rightarrow T$ and $C \rightarrow A$ ratios observed in Kap København.

**Table S1: Observed-to-expected ratios using UNREST vs GTR of *Betula*, *Salix*, *Populus*, and *Dryas* nuclear assigned reads from the Kap København samples.** Observed-to-expected ratios are scaled such that the smallest ratio is 1.000.

| | *Betula* | | *Salix* | | *Populus* | | *Dryas* | |
|---|---|---|---|---|---|---|---|---|
| Substitution | UNREST | GTR | UNREST | GTR | UNREST | GTR | UNREST | GTR |
| $A \rightarrow C$ | 1.0000 | 1.0000 | 1.2075 | 1.0598 | 1.0344 | 1.0215 | 1.0000 | 1.0000 |
| $C \rightarrow A$ | 1.4394 | 1.5055 | 1.3295 | 1.3817 | 1.2514 | 1.3577 | 2.9194 | 1.9901 |
| $A \rightarrow G$ | 1.1571 | 1.2254 | 1.0076 | 1.0101 | 1.0397 | 1.0983 | 1.2946 | 1.0732 |
| $G \rightarrow A$ | 3.8184 | 3.7669 | 2.8108 | 2.5735 | 2.7406 | 2.7972 | 8.0670 | 6.6644 |
| $A \rightarrow T$ | 1.0253 | 1.0106 | 1.1586 | 1.0369 | 1.0093 | 1.0046 | 1.3449 | 1.0634 |
| $T \rightarrow A$ | 1.0133 | 1.0097 | 1.1402 | 1.0264 | 1.0000 | 1.0000 | 1.3414 | 1.0674 |
| $C \rightarrow G$ | 1.0513 | 1.1088 | 1.2169 | 1.2442 | 1.0901 | 1.1738 | 1.2267 | 1.0552 |
| $G \rightarrow C$ | 1.0473 | 1.1013 | 1.2186 | 1.2417 | 1.0882 | 1.1702 | 1.2192 | 1.0501 |
| $C \rightarrow T$ | 4.0574 | 3.9769 | 2.9136 | 2.6599 | 2.8808 | 2.9236 | 8.7527 | 7.1723 |
| $T \rightarrow C$ | 1.1470 | 1.2133 | 1.0000 | 1.0000 | 1.0334 | 1.0916 | 1.2706 | 1.0525 |
| $G \rightarrow T$ | 1.4509 | 1.5002 | 1.3367 | 1.3801 | 1.2510 | 1.3506 | 2.9842 | 2.0150 |
| $T \rightarrow G$ | 1.0176 | 1.0092 | 1.2145 | 1.0635 | 1.0445 | 1.0343 | 1.0366 | 1.0381 |

#### S2.2 Kap København Modern Data Filtering

Oxidative damage manifests as elevated $G \rightarrow T$ and $C \rightarrow A$ substitutions (Costello et al. 2013). Library preparation can also generate these substitutions, so as a precaution we verified that the signal we attribute to ancient oxidative damage is not a preparation artifact. Damage introduced during steps shared across all libraries should affect modern and ancient reads comparably. Instead, the unfiltered modern reads carried higher putative oxidation than the ancient reads, alongside deamination higher than expected (Table S2). This suggests some of the unfiltered modern reads were introduced during preparation or index-hopped from the ancient libraries, rather than co-recovered alongside the aeDNA.

We reasoned that taxa more abundant in the negative controls than in the sediment samples are likely laboratory-derived, and filtered out any taxon with a higher read-per-sample rate in the controls than in the sediment samples. This reduced the modern set from 39 taxa (260,543 reads) to 4 taxa (5,404 reads). Both the deamination and oxidation ratios dropped after filtering. The filtered-out reads retained the elevated signal, whereas the remaining modern reads did not. The ancient reads remained greatly elevated throughout (Table S2) At 5,404 reads these estimates are noisy, with the $A \leftrightarrow T$ and $C \leftrightarrow G$ ratios exceeding the 1.0 expected under symmetry.

Laboratory and preparation processes are known to cause elevated $G \rightarrow T$, but the filtering shows this is not from a source shared between the modern and ancient reads. The filtered-out reads carried elevated $G \rightarrow T$ while the retained reads did not, so the ancient signal cannot be a preparation artifact common to all libraries.

**Table S2: Comparison of symmetric substitution ratios of ancient reads, modern filtered reads, and modern unfiltered reads.** Ratios are computed as $\max(i \rightarrow j, j \rightarrow i)/\min(i \rightarrow j, j \rightarrow i)$, so all values are $\geq 1.0$; values near 1.0 indicate symmetry.

| Filtering | Reads | $C \leftrightarrow T$ | $G \leftrightarrow A$ | $C \leftrightarrow A$ | $G \leftrightarrow T$ | $A \leftrightarrow T$ | $C \leftrightarrow G$ |
|---|---|---|---|---|---|---|---|
| Ancient | 192 258 396 | 5.7402 | 5.4297 | 1.9785 | 1.9239 | 1.0037 | 1.0021 |
| Unfiltered Modern | 260 543 | 1.6187 | 1.5964 | 2.4902 | 2.5092 | 1.0242 | 1.0045 |
| Filtered Modern | 5404 | 1.2510 | 1.2958 | 1.1213 | 1.1009 | 1.0732 | 1.1379 |

#### S2.3 Reference Databases

##### S2.3.1 Competitive Mapping Database

Reference database “WGS” consists of all NCBI (O’Leary et al. 2016) genomes filtered for ”reference genomes” underneath the Viridiplantae node, except the mistletoe genome (because it was extremely large), yielding 2298 total plant reference genomes. Additionally, we included those reference genomes underneath the NCBI taxonomic node IDs: 72037 6658 2864 3027 38254 2830 2570567 29197 2696291, which added a total of 213 references genomes across aquatic zooplankton, invertebrates and algae. Both databases were cut into chunks of approximately 20GB each, by either splitting using fasta-splitter 0.2.7 (Kryukov 2024) or concatenating genomes as needed (e.g. in the case of GTDB).

Eukaryotic:

- RefSeq’s mammalian vertebrates (vert_mam), plastid, mito and protozoas, downloaded 29/10/2024.
- PhyloNorway (Wang et al. 2021).
- Custom whole genome plant (and other taxa) sequences (”WGS”), downloaded from NCBI Genomes on 30/01/2025.

Microbial:

- GTDB v226.0 representative genomes (Parks et al. 2026)

##### S2.3.2 Krestovka Mammoth

The following nuclear and mitochondrial genomes were downloaded from NCBI RefSeq to build the Bowtie 2 (Langmead and Salzberg 2012) database to extract *Mammuthus primigenius.* This database is not comprehensive enough to taxonomically assign reads for molecular dating with ratePlacer, but enough to understand the pattern of DNA damage present.

- *Elephas maximus* nuclear genome: GCF_024166365.1
- *Mammuthus primigenius* mitochondrial genome: NC_007596
- *Homo sapiens* genome: GCF_000001405.40 (gh38)

##### S2.3.3 Kap København ratePlacer

For each of the following taxa (*Betula*, *Dryas*, *Populus*, and *Salix*), smaller databases were made for the compartment mapping with the nuclear genomes being used for ratePlacer. References were downloaded from RefSeq, GenBank (Clark et al. 2016), and Phytozome (Goodstein et al. 2012) and are listed in Table S3.

**Table S3: Reference IDs by genus, genetic compartment, and database.**

| Genus | Compartment | Database | Reference IDs |
|---|---|---|---|
| *Betula* | Chloroplast | RefSeq | NC_047177, NC_037473, NC_064121, NC_064120, NC_064122, NC_058835, NC_039992, NC_068733, NC_057498, NC_033978, NC_039993, NC_064119, NC_072281, NC_039994, NC_039995, NC_039996, NC_069292 |
| *Betula* | Mitochondria and Nuclear | GenBank | GCA_965285875 |
| *Betula* | Nuclear | Phytozome | *Betula platyphylla* v1.1 |
| *Alnus* | Chloroplast | RefSeq | NC_036751, NC_040996, NC_061930, NC_039930, NC_036752, NC_036753, NC_036754, NC_039991, NC_036755, NC_036756, NC_036757, NC_036758 |
| *Alnus* | Mitochondria and Nuclear | GenBank | GCA_965212435, GCA_958979055, GCA_965122795 |
| *Dryas* | Chloroplast | RefSeq | NC_088483, NC_088514 |
| *Dryas* | Chloroplast | GenBank | OY992843 |
| *Dryas* | Mitochondria and Nuclear | GenBank | GCA_963921425 |
| *Dryas* | Nuclear | GenBank | GCA_003254865, GCA_028570905, GCA_026122645 |
| *Purshia* | Chloroplast | RefSeq | NC_088513 |
| *Purshia* | Nuclear | GenBank | GCA_003254885 |

*continued on next page*

Table S3 – continued

| Genus | Compartment | Database | Reference IDs |
|---|---|---|---|
| *Populus* | Chloroplast | RefSeq | NC_008235, NC_009143, NC_024734, NC_024735, NC_024747, NC_027425, NC_028504, NC_031371, NC_031398, NC_032368, NC_032717, NC_033876, NC_036040, NC_037223, NC_037413, NC_037414, NC_037415, NC_037416, NC_037417, NC_037418, NC_037419, NC_037420, NC_037421, NC_040866, NC_040867, NC_040868, NC_040869, NC_040870, NC_040871, NC_040872, NC_040873, NC_040874, NC_040928, NC_040929, NC_040953, NC_044462, NC_045396, NC_047300, NC_058277, NC_058278, NC_058279, NC_058847, NC_060732, NC_080315 |
| *Populus* | Mitochondria | RefSeq | NC_028096, NC_028329, NC_035157, NC_041085 |
| *Populus* | Nuclear | GenBank | GCA_033621325, GCA_044906095, GCA_051312755, GCA_052148935, GCA_052724285, GCF_005239225 |
| *Salix* | Chloroplast | RefSeq | NC_024681, NC_026462, NC_026722, NC_028350, NC_035743, NC_035744, NC_036718, NC_037422, NC_037423, NC_037424, NC_037425, NC_037426, NC_037427, NC_037428, NC_037429, NC_043878, NC_044419, NC_051969, NC_053549, NC_054198, NC_056250, NC_056251, NC_056252, NC_056253, NC_057289, NC_057535, NC_058001, NC_058984, NC_058985, NC_058986, NC_059039, NC_060294, NC_060436, NC_061939, NC_063129, NC_063506, NC_063507, NC_064995, NC_068257, NC_068758, NC_068759, NC_069592 |
| *Salix* | Mitochondria | RefSeq | NC_029317, NC_029693, NC_046754, NC_052708, NC_052709, NC_058733, NC_064688, NC_068760, NC_068761, NC_069586, NC_088094 |
| *Salix* | Mitochondria and Nuclear | GenBank | GCA_040801835 GCA_040801865 GCA_964035475 |
| *Salix* | Nuclear | GenBank | GCA_049639585 |

For *Betula*, the selected outgroup was *Alnus*. For the compartment mapping, 17 *Betula* and 22 *Alnus* chloroplast, 2 *Betula* and 3 *Alnus* mitochondria, and 2 *Betula* and 3 *Alnus* nuclear genomes were downloaded. *Betula pendula* was used as the reference genome for `hal2maf` conversion (Armstrong et al. 2020).

For *Dryas*, the selected outgroup was *Purshia*. For the compartment mapping, 3 *Dryas* and 1 *Purshia* chloroplast, 1 *Dryas* and 0 *Purshia* mitochondria, and 4 *Dryas* and 1 *Purshia* nuclear genomes were downloaded. *Dryas octopetala* was used as the reference genome for `hal2maf` conversion.

*Populus* and *Salix* are outgroups of each other. For the compartment mapping, 44 *Populus* and 42 *Salix* chloroplast, 4 *Populus* and 15 *Salix* mitochondria, and 6 *Populus* and 4 *Salix* nuclear genomes were downloaded. For the *Populus* dating pipeline, *Populus deltoides* was used as the reference genome for `hal2maf` conversion while *Salix dunnii* was used for the *Salix* ratePlacer database.

#### S2.4 Genomic Compartment Filtering

A read that maps to more than one location may be placed in a region it did not originate from as nuclear copies of organellar DNA (NUMTs and NUPTs), transposable elements, or other duplicated and repetitive regions can recruit reads that belong elsewhere during mapping. A misplaced read carries the substitutions of its true origin into a position where they do not belong, and because these regions can be under different evolutionary constraints than the compartment they are placed in, the mismatch is not random. Since ratePlacer relies on the substitutions observed at each position, such mismapping would create discordance in the estimated age.

We retained a read only if it did not multimap within a single reference and did not multimap across genetic compartments (mitochondrial, nuclear, chloroplast). Reads that mapped uniquely with respect to both criteria were kept even if they multimapped across references within the same compartment, since our dataset contains multiple species and a read shared among them does not indicate a compartment or repeat-driven placement error. This removes reads whose genomic origin is ambiguous while retaining cross-species ambiguity that is expected given the reference set.

#### S2.5 Antarctic Subglacial Filtering and Data Extraction

The genera, along with their classification type (ancient, modern, or modern + surface), were taken from Figure 3 in De Sanctis et al. (2025) and are listed in Table **??**.

We cannot perform modern versus ancient analyses on the Antarctic samples, as the modern taxa reflect microbes that colonized the precipitate after it reached the surface, potentially over tens of thousands of years. Table S5 compares substitution ratios between subglacial and surface taxa along with the surface control. $C \rightarrow T$ remains elevated in both, consistent with deamination damage, but decreases on the surface control. $G \rightarrow T$ increases in the surface sample and surface control relative to the subglacial sample, possibly indicating that the surface environment promotes oxidative damage relative to the subglacial environment.

**Table S4: Antarctic microbial genera along with their IDs and environment classification.**

| Genus | Genus ID | Type |
|---|---|---|
| JAHJSF01 | 782751 | Ancient |
| TCS52 | 651589 | Ancient |
| Planktophila | 495357 | Ancient |
| Palsa-1315 | 483968 | Ancient |
| Desulfobacula | 502942 | Ancient |
| Nitrosarchaeum | 7005 | Ancient |
| Methanoperedens | 8412 | Ancient |
| JADFUQ01 | 717021 | Ancient |
| Polaromonas | 517395 | Ancient |
| Nitrotoga | 429612 | Ancient |
| Desulfatibia_A | 706597 | Ancient |
| UBA12170 | 785290 | Ancient |
| JAHJUB01 | 709076 | Ancient |
| 34-128 | 648504 | Ancient |
| Lutibacter | 632673 | Ancient |
| SPCO01 | 576161 | Ancient |
| SURF-13 | 632281 | Ancient |
| VSJD01 | 591753 | Ancient |
| RBG-16-66-20 | 545715 | Ancient |
| UBA2279 | 707064 | Ancient |
| 12-FULL-67-14b | 533847 | Ancient |
| JACCUC01 | 647858 | Modern + Surface |
| JACCVL01 | 598635 | Modern + Surface |
| SCTD01 | 675404 | Modern + Surface |
| CADCVQ01 | 630123 | Modern + Surface |
| JACCZJ01 | 735160 | Modern + Surface |
| Nocardioides | 416408 | Modern + Surface |
| JACDDK01 | 657643 | Modern + Surface |
| Aliterella | 668149 | Modern + Surface |
| Rubrobacter_D | 620438 | Modern + Surface |
| JAAYBF01 | 645787 | Modern + Surface |
| Pseudomonas_E | 212923 | Modern + Surface |
| JACDHN01 | 596575 | Modern + Surface |
| Corynebacterium | 275609 | Modern + Surface |
| Ramlibacter | 621309 | Modern |
| Arthrobacter_D | 589784 | Modern |
| Sphingorhabdus_B | 483518 | Modern |

**Table S5: Antarctic symmetry ratios for both subglacial and surface.** Ratios are computed as $\max(i \to j, j \to i)/\min(i \to j, j \to i)$, so all values are $\geq 1.0$; values near 1.0 indicate symmetry.

| Sample | Library | $C \leftrightarrow T$ | $G \leftrightarrow A$ | $C \leftrightarrow A$ | $G \leftrightarrow T$ | $A \leftrightarrow T$ | $C \leftrightarrow G$ |
|---|---|---|---|---|---|---|---|
| Antarctic (subglacial) | ss | 2.7875 | 1.1133 | 1.0781 | 1.0939 | 1.0440 | 1.0339 |
| Antarctic (surface) | ss | 2.0560 | 1.1197 | 1.1577 | 1.2721 | 1.0032 | 1.0695 |
| Antarctic (surface control) | ss | 1.4776 | 1.1857 | 1.2885 | 1.5179 | 1.0322 | 1.0802 |
